# Biological Aging in Childhood and Adolescence Following Experiences of Threat and Deprivation: A Systematic Review and Meta-Analysis

**DOI:** 10.1101/642405

**Authors:** N.L. Colich, M.L. Rosen, E.S. Williams, K.A. McLaughlin

## Abstract

Life history theory argues that exposure to early-life adversity (ELA) accelerates development, although existing evidence for this varies. We present a meta-analysis and systematic review testing the hypothesis that ELA involving threat (e.g., violence exposure) will be associated with accelerated biological aging across multiple metrics, whereas exposure to deprivation (e.g., neglect, institutional rearing) and low-socioeconomic status (SES) will not. We meta-analyze 46 studies (n=64,925) examining associations of ELA with pubertal timing and cellular aging (telomere length and DNA methylation age), systematically review 19 studies (n=2276) examining ELA and neural markers of accelerated development (cortical thickness and amygdala-prefrontal cortex functional connectivity) and evaluate whether associations of ELA with biological aging vary according to the nature of adversity experienced. ELA overall was associated with accelerated pubertal timing (d=-0.12) and cellular aging (d=-0.32). Moderator analysis revealed that ELA characterized by threat (d=-0.26), but not deprivation or SES, was associated with accelerated pubertal development. Similarly, exposure to threat-related ELA was associated with accelerated cellular aging (d=-0.43), but not deprivation or SES. Systematic review revealed associations between ELA and accelerated cortical thinning, with threat-related ELA consistently associated with thinning in ventromedial prefrontal cortex, and deprivation and SES associated with thinning in frontoparietal, default, and visual networks. There was no consistent association of ELA with amygdala-PFC connectivity. These findings suggest specificity in the types of early environmental experiences associated with accelerated biological aging and highlight the importance of evaluating how accelerated aging contributes to health disparities and whether this process can be mitigated through early intervention.

## Introduction

Exposure to early-life adversity (ELA)—including exposure to child abuse, sexual assault, neglect, and chronic poverty—is associated with elevated risk for numerous mental and physical health problems, including depression, anxiety disorders, substance abuse, suicide, and cardiovascular disease (Felitti et al., 1998; Green et al., 2010; Heim & Binder, 2012; Kessler et al., 2010; McLaughlin et al., 2010, 2012; Norman et al., 2012; Scott et al., 2011). The associations of ELA with mental and physical health problems are observable beginning in childhood and adolescence (Boynton-Jarrett, Ryan, Berkman, & Wright, 2008; Halpern et al., 2013; McLaughlin et al., 2012; McLaughlin, Basu, et al., 2016) and persist into adulthood (Dong et al., 2004; Felitti et al., 1998; Green et al., 2010; Kessler et al., 2010). Recent evidence from longitudinal and population-based studies indicates that exposure to ELA is even associated with elevated risk for premature mortality (Brown et al., 2009; E. Chen, Turiano, Mroczek, & Miller, 2016).

### Accelerated Development / Biological Aging

One potential mechanism linking exposure to ELA with this wide range of physical and mental health problems is accelerated biological aging. Specifically, exposure to adversity early in life may alter the pace of development, resulting in faster aging. Most conceptual models on the link between ELA and accelerated development are based in Life History Theory (J. Belsky, Steinberg, & Draper, 1991; Ellis, Figueredo, Brumbach, & Schlomer, 2009; Ellis & Garber, 2000), which postulates that experiences in early-life can program an individual’s developmental trajectory in order to respond most effectively to the environmental demands they are likely to encounter later in life. The pattern and timing of life history events—such as age of sexual maturation, gestational period, number of offspring, birth spacing, length of parental investment, longevity, and others—is determined by the relative prioritization of time and energy invested in growth, reproduction, and longevity (Del Giudice, Gangestad, & Kaplan, 2016; Hill & Kaplan, 1999). For instance, in a safe, predictable and enriched environment, a slow and protracted development may be optimal, as it allows for maximal parental investment prior to offspring independence. However, in a harsh or unpredictable environment, a faster pace of development in which individuals reach adult-like capabilities at an earlier age may be favored in order to maximize reproduction prior to potential mortality. Life history theories of human development argue that early environments characterized by harshness (e.g. trauma, violence exposure) may accelerate the onset of puberty in order to maximize the opportunity for reproduction prior to mortality (J. Belsky, 2012; Ellis et al., 2009; Rickard, Frankenhuis, & Nettle, 2014). However, in unpredictable environments, where there is large variation in harshness, it may be optimal to delay reproductive milestones, depending upon various features of the environment including population density and resource availability (J. Belsky, 2012; Ellis et al., 2009).

More recently, life history theories regarding the pace of development following ELA have been extended to focus on additional measures of biological aging. First, predictive adaptive response models (Nettle, Frankenhuis, & Rickard, 2013; Rickard et al., 2014) focus on cellular and molecular development and how it relates to an individual’s morbidity and mortality across the lifespan. These models propose that ELA negatively influences physical health, through altered cellular development as a result of reduced energy to build or repair cellular tissue. This advanced cellular aging forecast may reduce longevity and contribute to acceleration in reproductive maturity (Nettle et al., 2013; Rickard et al., 2014). Second, the stress acceleration hypothesis (Callaghan & Tottenham, 2016) suggests that ELA accelerates development of neural circuitry underlying emotional processing, specifically, development of the amygdala-prefrontal cortex (PFC) circuit thought to underlie emotion regulation capabilities. This accelerated development in the context of unreliable or absent caregiving may occur in order to allow for independent emotion regulation at an earlier age (Callaghan & Tottenham, 2016). Each of these theories rest on the assumption that ELA impacts the pace of development across multiple domains and metrics of biological aging.

### ELA and Biological Aging

Biological aging following ELA has been measured with a variety of different metrics. By far the most commonly used metric is the timing and pace of pubertal development, including age of menarche in females (Boynton-Jarrett & Harville, 2012; Deardorff, Abrams, Ekwaru, & Rehkopf, 2014; Graber, Brooks-Gunn, & Warren, 1995b) and pubertal stage controlling for chronological age (Colich et al., 2019; Mendle, Leve, Van Ryzin, Natsuaki, & Ge, 2011; Negriff, Blankson, & Trickett, 2015; Noll et al., 2017; Sumner, Colich, Uddin, Armstrong, & McLaughlin, 2019). A second line of work has examined measures of cellular aging, including leukocyte telomere length (Coimbra, Carvalho, Moretti, Mello, & Belangero, 2017; Drury et al., 2014; Price, Kao, Burgers, Carpenter, & Tyrka, 2013) and DNA methylation (DNAm) age (Gassen, Chrousos, Binder, & Zannas, 2017; Wolf et al., 2017). A separate literature has examined markers of neural maturation such as amygdala-PFC connectivity (Callaghan & Tottenham, 2016; Gee, Gabard-Durnam, et al., 2013) and cortical thickness (McLaughlin, Sheridan, Winter, et al., 2014).

Evidence for accelerated biological aging following ELA has been found across all of these metrics. For example, numerous studies have found that ELA is associated with earlier pubertal timing (Graber, Brooks-Gunn, & Warren, 1995a; Hartman, Li, Nettle, & Belsky, 2017; Mendle et al., 2011; Negriff, Blankson, et al., 2015). Similarly, a small but increasing number of studies have reported accelerated cellular aging following ELA, including shorter telomere length (Drury et al., 2012, 2014; Mitchell et al., 2014; Shalev et al., 2013), and advanced DNAm age relative to chronological age (Jovanovic et al., 2017a; Sumner et al., 2019). Finally, much of the evidence for accelerated neural development following ELA comes from studies examining amygdala-PFC functional connectivity (Colich et al., 2017; Gee, Gabard-Durnam, et al., 2013; Keding & Herringa, 2016) and cortical thinning across development (McLaughlin, Sheridan, Winter, et al., 2014). However, other studies have found no associations between ELA and pubertal timing (Negriff, Saxbe, & Trickett, 2015; Negriff & Trickett, 2012) or cortical thinning (McLaughlin, Sheridan, et al., 2016; Rosen, Sheridan, Sambrook, Meltzoff, & McLaughlin, 2018). Some studies have even found that ELA is associated with slower or delayed pubertal timing (Johnson et al., 2018; Negriff, Blankson, et al., 2015; Sumner et al., 2019) and a more immature pattern of amygdala-PFC connectivity (Cisler, James, et al., 2013; Marusak, Martin, Etkin, & Thomason, 2015; Silvers, Lumian, et al., 2016). The strength and direction of the association between ELA and markers of biological aging varies widely across studies, and to date no systematic review or meta-analysis on this topic has been conducted.

We argue, and test through meta-analysis and systematic review, that the wide variability in the association of ELA with accelerated development might be explained—at least in part—by differences in how distinct types of ELA influence the pace of development. Existing studies have focused on a wide range of adversity experiences, ranging from physical abuse and violence exposure to physical and emotional neglect and institutional rearing, and provide some clues about the types of ELA that might be particularly likely to produce a pattern of accelerated development. For example, physical and sexual abuse have been consistently associated with accelerated pubertal development in females (Mendle, Ryan, & McKone, 2016; Natsuaki, Leve, & Mendle, 2011; Noll et al., 2017; Trickett, Noll, & Putnam, 2011; Trickett & Putnam, 1993). In contrast, studies of war and famine suggest that severe material deprivation can delay pubertal development (Prebeg & Bralic, 2000; van Noord & Kaaks, 1991). Although less work has examined the effects of neglect and psychosocial deprivation on biological aging, existing studies typically find no association of neglect or early institutional rearing with pubertal timing (Johnson et al., 2018; Mendle et al., 2011; Reid et al., 2017; Ryan, Mendle, & Markowitz, 2015). In contrast, early institutionalization is associated with an accelerated pattern of cellular aging (Drury et al., 2012) and maturation of the amygdala-PFC circuit (Gee, Gabard-Durnam, et al., 2013), suggesting that accelerated biological aging might not occur in a uniform manner across various neurobiological systems.

Discrepancies in these findings may be due to the treatment of ELA as a monolithic construct with equifinality across all metrics of biological aging. To date, no attempt has been made to consider how associations of ELA with accelerated biological aging might vary according to the nature of the adversity experienced. Systematic investigation into variability in the association of ELA with biological aging across adversity types may help to reconcile inconsistent findings and advance theoretical models of how early experiences alter the pace of development at reproductive, cellular, and neural levels of analysis. This meta-analysis aims to do so by: 1) examining how *differing dimensions* of ELA influence biological aging, distinguishing between experiences characterized by threat versus deprivation; and 2) evaluating whether the associations of these different types of adverse early experience with biological aging are global or specific to particular domains of aging—including pubertal timing, cellular aging and brain development.

### Conceptual Model of Early-Life Adversity and Accelerated Development

Many prior studies examining the effects of ELA on accelerated biological aging have focused on a limited range of ELA experiences, typically focusing on relative extreme exposures like sexual abuse or institutional rearing. Other studies have utilized a cumulative-risk approach, which tallies the number of distinct forms of ELA experienced to create a risk score without regard to the type, chronicity, or severity of the experience and uses this risk score as a predictor of outcomes, with the assumption that all forms of ELA have equal and additive effects on developmental outcomes (Evans, Li, & Whipple, 2013). Very few studies attempt to address the high co-occurrence of varying forms of ELA (Green et al., 2010; McLaughlin et al., 2012) or examine the differential influences of particular adversity types on biological aging, with some notable exceptions (Mendle et al., 2011, 2016; Negriff, Saxbe, et al., 2015; Sumner et al., 2019).

The dimensional model of adversity and psychopathology (DMAP) argues that the wide range of experiences currently classified as ELA can be organized into core underlying dimensions that have unique influences on cognitive, emotional, and neural development (McLaughlin, Sheridan, & Lambert, 2014; McLaughlin & Sheridan, 2016; Sheridan & McLaughlin, 2014). This model attempts to distill complex adverse experiences into core underlying dimensions that cut across multiple forms of ELA that share common features. Two such dimensions are *threat*, which encompasses experiences involving harm or threat of harm to the child, and *deprivation*, which involves an absence of expected inputs from the environment during development, such as cognitive and social stimulation (e.g., complex language directed at the child) as well as emotional nurturance (e.g., emotional neglect). In addition, the DMAP model argues that these dimensions of adversity have influences on emotional, cognitive, and neural development that are at least partially distinct. Increasing evidence has demonstrated the unique developmental consequences of threat and deprivation on developmental outcomes (Busso, McLaughlin, & Sheridan, 2016; Dennison et al., 2019; Everaerd et al., 2016; Lambert, King, Monahan, & McLaughlin, 2017; Rosen et al., 2018; Sheridan, Peverill, Finn, & McLaughlin, 2017). Determining whether all forms of ELA are associated with accelerated development across multiple metrics of biological aging or whether only particular dimensions of ELA are associated with this pattern is critical for identifying the mechanisms linking ELA to health outcomes and to better inform early interventions.

The threat dimension of ELA is conceptually similar to the life history theory dimension of environmental harshness, and involves experiences of trauma and violence exposure. We expect that experiences characterized by threat will be associated with accelerated biological aging, potentially in order to maximize the opportunity for reproduction prior to mortality (J. Belsky, Schlomer, & Ellis, 2012; Ellis et al., 2009). However, it is unclear how experiences of deprivation align with life history theory; whereas nutritional deprivation and food insecurity are thought to delay pubertal timing to ensure maximal bioenergetic resources should reproduction occur (Rogol, Clark, & Roemmich, 2000), specific predictions about physical and emotional neglect are lacking in life history models. Preliminary evidence suggests that accelerated development following ELA may vary across different dimensions of adversity. For instance, we have found that experiences characterized by threat, but not deprivation, were associated with accelerated pubertal stage relative to chronological age and accelerated DNAm age in a community-based sample of children and adolescents (Sumner et al., 2019). In contrast, experiences characterized by deprivation were associated with *delayed* pubertal timing, after controlling for co-occurring threat experiences. We recently replicated this work in a nationally-representative sample of adolescent females, using age of menarche as our metric of accelerated aging (Colich et al., 2019). Determining whether accelerated biological aging is associated with exposure to ELA generally or with particular dimensions of ELA may help to elucidate the specific psychological and biological mechanisms underlying these associations.

### Metrics of Biological Aging

Accelerated biological aging has been conceptualized in many ways, across multiple domains of biological development. Historically, these domains have been examined in isolation, independent of other domains of biological aging. Only two studies to our knowledge have explored the effects of ELA on multiple domains of accelerated development in adolescence (J. Belsky & Shalev, 2016; Sumner et al., 2019) and recent work suggests that accelerated telomere erosion and accelerated pubertal development represent similar biological processes as a consequence of ELA (Shalev & Belsky, 2016).

#### Pubertal timing

The most consistently examined marker of accelerated development in relation to ELA is pubertal timing, typically operationalized as the age of onset of pubertal development, or the age of achieving a reproductive milestone such as menarche. Puberty begins as early as ages 8-14 in females and 9-15 in males with the activation of the hypothalamic-pituitary-gonadal (HPG) axis. This ultimately initiates the start of gonadarche, in which the gonads mature and produce gondadal hormones or sex steroids. This in turn, leads to breast development and eventually menarche in girls, and in increased testicle size and the onset of spermarche in males. Typical measures of pubertal development use secondary sex characteristics as a metric of pubertal stage (Carskadon & Acebo, 1993; Marshall & Tanner, 1969, 1970). For the purposes of this meta-analysis, we have included studies that explore how ELA is associated with three commonly used metrics of pubertal timing—pubertal stage relative to chronological age, age at the achievement of the onset of secondary sex characteristics, and age of menarche. Although menarche occurs relatively late in the pubertal process, participants are relatively reliable in their reporting of this milestone, particularly in adolescence (Dorn, Sontag-Padilla, Pabst, Tissot, & Susman, 2013).

#### Cellular aging

Predictive adaptive response models of accelerated aging following adversity postulate that the early environment influences an individual’s somatic state, which in turn influences reproductive timing and other life history events (Nettle et al., 2013; Rickard et al., 2014). One pathway linking ELA to somatic states is cellular aging. Some argue that if the body detects a shortened cellular lifespan, mechanisms may exist to accelerate the development of the reproductive system in order to maximize the chances of reproduction prior to mortality (Nettle et al., 2013; Rickard et al., 2014). Cellular aging in the context of ELA has been measured in two different ways – telomere length and metrics of epigenetic aging using DNAm patterns.

Telomeres are nucleopeptide complexes that sit at the end of chromosomes and protect the chromosome from degradation (Chan & Blackburn, 2004). Telomeres shorten due to both cell replication and exposure to oxidative stress and inflammation. In normal aging, telomeres shorten in all cell types, which allows for the use of telomere length as a biological marker of cellular age (Frenck, Blackburn, & Shannon, 1998). Chronic stress has been shown to accelerate the shortening of telomeres in adults (Epel et al., 2004), and several studies have demonstrated associations between ELA and telomere length in children (Coimbra et al., 2017; Essex et al., 2013; Price et al., 2013). Shortened telomere length has been implicated in the pathogenesis of both physical and mental health problems in adulthood (Gotlib et al., 2015; Hoen et al., 2013; Needham, Mezuk, et al., 2015; Tyrka et al., 2016), suggesting a potential mechanism linking ELA and maladaptive health outcomes in adolescence and adulthood.

A second recently established metric of cellular aging is an epigenetic clock that considers genome-wide DNAm patterns (both increased and decreased methylation of select CpG sites) to quantify biological age independent from chronological age (DNAm age) (Hannum et al., 2013; Horvath, 2013). This metric correlates strongly with chronological age in both adolescents and adults (Horvath & Raj, 2018; Suarez et al., 2018) and shows strong positive associations with age of death (B. H. Chen et al., 2016; Marioni, Shah, McRae, Chen, et al., 2015), suggesting it is a valid metric of cellular aging. Deviations between DNAm age and chronological age have been used as a metric of accelerated development (Davis et al., 2017; Jovanovic et al., 2017b; Sumner et al., 2019) and are associated with exposure to ELA (Jovanovic et al., 2017b; Sumner et al., 2019). Advanced DNAm age has been associated with increased risk of cardiovascular disease, cancer, and obesity (Horvath et al., 2014; Marioni, Shah, McRae, Ritchie, et al., 2015; Perna et al., 2016), again potentially highlighting a mechanism linking ELA and physical health problems.

For the purpose of this meta-analysis, we have included studies that explore how ELA impacts cellular aging, as measured by telomere length and DNAm age.

#### Brain Development

Numerous studies have investigated the neural consequences of ELA. Here, we focus specifically on neural markers of maturation. As such, we focus on two metrics for which patterns of development have been well characterized: cortical thickness and functional connectivity between the amygdala and prefrontal cortex (PFC). We focus on cortical thickness as a metric of structural development because the pattern of development is well characterized, replicated across many studies, and shows a clear linear association with age, such that cortical thickness steadily decreases from middle childhood to early adulthood (Ducharme et al., 2016; LeWinn, Sheridan, Keyes, Hamilton, & McLaughlin, 2017; Vijayakumar et al., 2016; Wierenga, Langen, Oranje, & Durston, 2014). Second, we focus on functional connectivity between the amygdala and PFC as a metric of maturation because it serves as a key component in the stress acceleration hypothesis, which posits that the amygdala-PFC circuit supporting emotional processing and regulation matures more rapidly among children exposed to ELA (Callaghan & Tottenham, 2016).

Cortical thickness declines steadily from childhood to early adulthood (Ducharme et al., 2016; LeWinn et al., 2017; Wierenga et al., 2014), as a result of developmentally-appropriate pruning of synapses and increases in myelination of connections between neurons (Natu et al., 2018; Sowell et al., 2004). This linear pattern of development enables assessment of whether development is accelerated or delayed among children with ELA relative to their peers. Cortical structure can be measured in a variety of ways including surface area, thickness, and volume (for review, see Vijayakumar et al., 2016). However, cortical thickness is the only metric that has a linear developmental trajectory, declining steadily from early childhood through early adulthood (Ducharme et al., 2016; LeWinn et al., 2017; Walhovd, Fjell, Giedd, Dale, & Brown, 2017; Wierenga et al., 2014). In contrast, cortical surface area and volume exhibit non-linear associations with age and the inflection points of these trajectories vary across samples and remain a source of debate (Ducharme et al., 2016; Giedd et al., 1999; Lenroot et al., 2007; LeWinn et al., 2017; Mills et al., 2016; Vijayakumar et al., 2016). These non-linear patterns of development make assessing deviations from the expected pattern more difficult. Therefore, we focus only on studies that use cortical thickness—including both whole cortex and specific regions—as an outcome.

The stress acceleration hypothesis focuses on the impact of ELA on the developmental trajectory of neural circuits supporting emotion processing and regulation, particularly on connectivity between the amygdala and medial PFC (mPFC; (Callaghan & Tottenham, 2016). Animal tracing studies demonstrate that feedforward connections between amygdala and PFC exist early in life, but feedback connections emerge later in development (Barbas & García-Cabezas, 2016). It has been proposed that in humans, changes in functional connectivity between the mPFC and amygdala may reflect the maturation of these feedback connections (Gee, Humphreys, et al., 2013). The pattern of functional connectivity between the amygdala and the mPFC shifts from positive to negative across development in the context of emotional processing tasks (Gee, Humphreys, et al., 2013; Silvers, Lumian, et al., 2016; Wu et al., 2016).

For the purpose of this systematic review, we have included studies that explore how ELA impacts both cortical thickness and amygdala-mPFC functional connectivity.

### The Current Study

We aimed to test the hypothesis that experiences characterized by threat, but not deprivation, would accelerate biological aging. Applying this theoretical framework may help to reconcile discrepant findings in the literature by evaluating how different dimensions of ELA influence biological aging. In addition, we aimed to integrate disparate literatures by examining whether different dimension of adversity have general or specific effects on multiple domains of biological aging—including pubertal timing, cellular aging and brain development. We expected that threat and deprivation will have differing effects on biological aging, with threat associated with accelerated biological aging across all metrics and deprivation associated with delayed pubertal development. We did not have specific hypotheses about how deprivation would influence cellular aging or brain development. We also separately examined the associations of socioeconomic status (SES) with biological aging, as SES is a commonly used global measure of early experience that is associated with increased risk of exposure to both threat and deprivation (e.g., Green et al., 2010; McLaughlin et al., 2012). We had no a priori hypotheses about SES, given these associations with both threat and deprivation. A final guiding question was whether the associations of ELA characterized by threat and deprivation with biological aging would be consistent across all metrics. Whereas pubertal development reflects a more global measure of aging, cellular aging is a metric of biological aging most relevant to physical health, and cortical thickness and development in the amygdala-PFC circuit may reflect learning or adaptation to a stressful early environment, but not aging in a global way.

## Methods

### Information Sources and Search Strategy

This meta-analysis and systematic review was conducted in line with the PRISMA guidelines for meta-analyses (Moher, Liberati, Tetzlaff, & Altman, 2009; Figure 1). To identify studies with relevant data, literature searches were conducted using internet databases (PubMed, SCOPUS, PsycINFO, Web of Science and Google Scholar) through January 2018. To ensure a thorough search, search terms encompassed various forms of ELA (e.g., violence, trauma, neglect, maltreatment, institutional rearing, deprivation, SES, poverty, early adversity, early life stress) as well as our dependent measures of interest (e.g., puberty, cell aging, methylation, menarche, telomere length, methylation, neural) and our targeted study population (e.g., infant, child, adolescent, pediatric) (see Supplemental Information for all search terms). All included studies were published in English and from peer-reviewed journals. To further identify eligible studies, we reviewed references of identified papers for additional studies using forward and backward searching.

**Figure 1.**
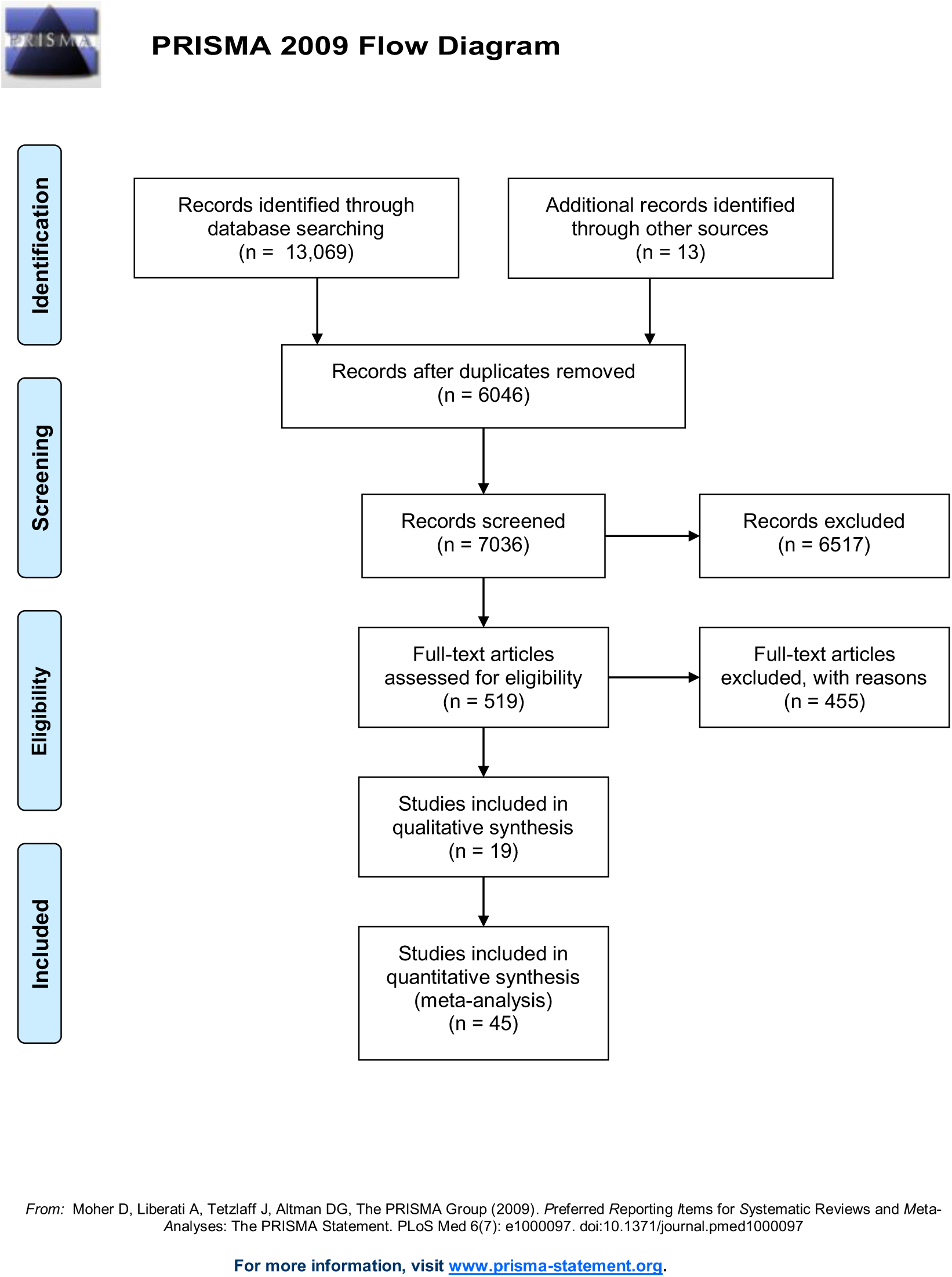
2009 PRISMA flow diagram of literature search and screening process.

#### Study inclusion criteria

In order to be included in the meta-analysis, studies had to meet the following criteria. First, studies had to examine an association between ELA and one of our dependent measures (pubertal timing, cellular aging, or brain development) and report sufficient statistics to calculate an effect size. Second, studies had to have been conducted in children and adolescent human samples (participants under age 18), rather than using retrospective reports of ELA and development in adults given the well documented recall biases associated with retrospective reporting of childhood experiences in adulthood (Hardt & Rutter, 2004; Widom, Raphael, & DuMont, 2004).

##### Inclusion criteria for ELA

We draw on a recent definition of ELA as experiences that were either chronic or severe in nature that require psychological or neurobiological adaptation by an average child and that represent a deviation from the expectable environment (McLaughlin, 2016). As detailed above, we used a wide range of search terms for ELA encompassing maltreatment experiences (e.g., physical, sexual, and emotional abuse; physical and emotional neglect), exposure to traumatic events (e.g., observing domestic violence, being the victim of interpersonal violence), institutional rearing, material deprivation (e.g., food insecurity), and childhood SES. We did not consider biological father absence as a form of ELA given that: a) it is not clearly a form of ELA based on prevailing definitions (McLaughlin, 2016); and b) a meta-analysis on father absence and pubertal timing was recently conducted (Webster, Graber, Gesselman, Crosier, & Schember, 2014). We did not include other early experiences or more global stressful life events that did not clearly meet our definition of ELA (e.g., parental psychopathology, peer victimization).

##### Inclusion criteria for studies of pubertal timing

To retain as many studies as possible, we included studies that used self-report, parent-report and physician-rated Tanner/PDS stage (controlling for age) or age of menarche. Physician-rated Tanner stage and interview-based assessments of age of menarche in adolescence have been shown to be acceptably reliable (Coleman & Coleman, 2002; Dorn & Biro, 2011; Dorn et al., 2013). We examined whether the specific measure of pubertal timing was a moderator of ELA-puberty associations.

##### Inclusion criteria for studies of cellular aging

Although there have been prior reviews and meta-analyses exploring the effects of ELA on telomere length (Coimbra et al., 2017; Price et al., 2013) or DNAm age (Gershon & High, 2015; Lewis & Olive, 2014; Silberman, Acosta, & Zorrilla Zubilete, 2016; Vinkers et al., 2015; Wolf et al., 2018), none has focused on differences across distinct adversity types or restricted the focus to studies measuring ELA and cellular aging in childhood or adolescence. Telomere length and DNAm age can be assessed through both blood and saliva samples (Wren, Shirtcliff, & Drury, 2015); we have included both measures in our analyses.

##### Inclusion criteria for studies of brain development

We included only studies that assessed cortical thickness—including both whole cortex and specific regions—as an outcome and not other measures of cortical structure (e.g., volume and surface area) where age-related patterns are non-linear and thus more difficult to interpret with regard to acceleration of development. If ELA-exposed youth exhibit thinner cortex than non-exposed youths of the same age, this will be interpreted as accelerated maturation; if ELA-exposed youths exhibit thicker cortex than non-exposed youths of the same age, this will be interpreted as delayed development. Similarly, we focus only on studies exploring task-related amygdala-mPFC functional connectivity, where a developmental shift from positive to negative in task-related amygdala-mPFC connectivity has been documented (Callaghan & Tottenham, 2016; Gee, Gabard-Durnam, et al., 2013). We will evaluate studies of ELA with this normative developmental pattern in mind; if children who have experienced adversity demonstrate greater negative connectivity for their age than children who have not, this would reflect accelerated development and if children who have experienced adversity exhibit more positive or less negative connectivity than comparison children, this would reflect delayed development. In contrast, studies investigating developmental patterns of connectivity at rest have been more mixed, with some studies demonstrating an increase in connectivity with age (e.g., Gabard-Durnam et al., 2014) and others demonstrating a decrease (Jalbrzikowski et al., 2017). Because a consensus has not been reached on the normative developmental pattern of amygdala-mPFC connectivity during rest, we focus only on papers that explore the associations of ELA with amygdala-mPFC connectivity using task-related functional connectivity.

We conducted a systematic review for metrics of brain development rather than a meta-analysis for the following reasons. First, whole-brain fMRI meta-analyses focus on the spatial nature of associations across the brain as opposed to the strength of effect sizes within a designated region. Given our focus on a specific measure of functional connectivity (amgydala-mPFC), spatial maps do not sufficiently address our research question regarding this metric of neural development. Second, the use of heterogenous ROIs in studies of cortical thickness and amygdala-mPFC connectivity (e.g., different regions of mPFC), make it difficult to quantitatively compare results across studies. Third, meaningful differences in task design and task demands make it difficult to directly compare results of amygdala-mPFC connectivity using meta-analysis.

### Measuring and Coding Adversity

In order to directly compare results from the present study with the majority of the existing literature, we first conducted an analysis in which we include studies defining ELA broadly, regardless of adversity type. We then examined whether associations of ELA with biological aging metrics exhibited significant heterogeneity, and evaluated whether adversity type (i.e., threat, deprivation, SES) was a moderator of these associations (see Analysis Methods for details). Consistent with previous work from our group (Colich et al., 2019; Dennison et al., 2019; McLaughlin, Sheridan, Lambert, et al., 2014; Sheridan & McLaughlin, 2014), we conceptualized threat-related adversities to include experiences of physical abuse, domestic violence, sexual assault, witnessing or being the victim of violence in the community, and emotional abuse. Deprivation-related adversities included physical neglect, low cognitive stimulation, food insecurity, and early institutionalization/international adoption. We also examined the effects of SES, including family income and parental education. Although low SES is associated with reductions in cognitive stimulation among children (Bradley, Corwyn, Burchinal, McAdoo, & Garcia Coll, 2001; Duncan & Magnuson, 2012; Gilkerson et al., 2017), SES is a proxy for deprivation rather than a direct measure. This is especially true when studies examine the effects of SES without controlling for co-occurring experiences of threat or violence. To ensure that we had not diluted our deprivation composite by including SES as an indicator, we chose to examine studies using SES as a metric of ELA separately.

#### Study exclusion criteria

The literature search yielded a total of 7081 studies. Studies were first excluded based on their title or abstract (k=6515) with exclusion decisions made by one of the authors (NLC, EAW or MLR) and confirmed by another. Exclusion criteria included any publication that was not an analysis of primary data (i.e. a review, book chapter, etc.). We also excluded any studies conducted outside of the United States, Western Europe or Australia, given well-documented effects of ethnicity, nutritional status, and SES on timing of development (Parent et al., 2003) and difficulties assessing SES consistently across different countries. A subset of studies were examined more thoroughly for eligibility (k=519). After a careful review of the methods, studies were excluded if they didn’t include a relevant independent or dependent variable (i.e. single gene methylation patterns rather than a measure of epigenetic aging or resting state amygdala-PFC connectivity rather than task-based connectivity; k=136), if they were a review paper or book chapter (k=134), if they were conducted outside of the US/Europe/Australia (k=81), if the study was conducted in infants, adults or nonhuman animals (k=56), if we were unable to access the manuscript (n=24), if the data were from a conference abstract (k=8), published in a foreign language only (k=3), if the study was not sufficiently powered (i.e. less than or equal to 5 participants per group; k=2), or the study was later retracted (k=1). Given our focus on understanding deviations in developmental timing following ELA, studies were also excluded if the exposed and control group differed significantly in age (k=1; Humphreys et al., 2016). Finally, studies were excluded if they did not include data that we were able to convert into an effect size after multiple attempts to contact the study authors for original data; (k=9). Overall, the current study included a total of 64 studies: 37 studies contributing to our meta-analysis exploring the effects of ELA on pubertal timing, 9 studies contributing to our meta-analysis exploring the effects of ELA on cellular aging, and 19 studies contributing to our systematic review exploring the effects of ELA on brain development (with one study [Sumner et al., 2019] contributing to both pubertal timing and cellular aging analyses.

### Management of Non-Independent Samples

In many cases, we extracted multiple effect sizes from the same sample. For example, some studies included multiple measures of pubertal timing (e.g. pubertal stage and age of menarche) or multiple measures of ELA (e.g. sexual abuse and physical abuse). Similarly, associations between ELA and developmental timing from a single study were sometimes examined across multiple publications using the same sample. To deal with this non-independence, we conducted multilevel mixed effects analyses with restricted maximum likelihood estimation, including study nested within sample as a random effect, such that multiple effect size estimates are nested within a higher-level grouping variable (e.g. study or sample). In the case of longitudinal data, we always included data from Wave I, as this wave tends to have the lowest attrition rate and in turn, the largest sample size (Borenstein, Hedges, Higgins, & Rothstein, 2009). If a separate paper included data from a later wave, we included that data and did not report data from Wave I a second time (as these associations were included in the analysis from the Wave I paper; for example, (Mendle, Leve, Van Ryzin, & Natsuaki, 2014; Mendle et al., 2011).

### Data Extraction

Three trained raters (NLC, MLR, EAW) coded individual studies. We screened each study and coded variables for study year, authors, participant composition/sample, mean age of participants, number of males and female participants, ethnicity, pubertal timing measure and informant, ELA measure and informant, as well as whether analyses controlled for other types of adversity, parent psychopathology, child psychopathology, father absence, mother’s age at menarche, and BMI. All disagreements in coding were resolved via discussion amongst the three raters until consensus was achieved.

### Data Analysis

To ensure consistency in the directionality of the effect sizes, in all cases, metrics of developmental timing were coded to indicate that numerically lower values (negative values) indicated accelerated development, to be consistent with age at menarche or age at pubertal attainment (the most commonly used metrics). Similarly, adversities were coded so that a numerically higher value indicates greater adversity. To be consistent with other variables, SES was coded to indicate that numerically higher SES values indicate lower SES.

For each study we calculated an effect size *d* and corresponding sampling variance (Cohen’s d; Cohen, 1988) for each relevant analysis. Following Cohen’s suggestion, we interpret a *d=*0.2 to be a ‘small’ effect, *d=*0.5 to be a ‘medium’ effect and *d=*0.8 to be a ‘large’ effect. A positive *d* indicates that exposure to ELA is associated with delayed development (later age at a developmental milestone), where as a negative *d* indicates that greater adversity is associated with accelerated development (earlier age at a developmental milestone). We derived *d*s from multiple reported statistics including: unadjusted or adjusted correlations between two variables, odds ratios, mean differences and standard deviations, *t* statistics, *F* statistics and associated *Ns* and *p* values, as well as unstandardized and standardized regression coefficients. All effect sizes were computed in R (version 3.4.1) using the escalc function in the “metafor” package (Viechtbauer, 2010) and converted to Cohen’s *d* using established formulas (Borenstein et al., 2009). Authors were contacted when published manuscripts met criteria for inclusion but did not include the necessary data to calculate an effect size amenable to our analyses, which occurred in thirteen cases (of which 4 provided necessary data and were included in our analyses).

All meta-analysis were conducted using three-level mixed-effects models and the rma.mv function in the “metafor” package (Viechtbauer, 2010) in R (version 3.4.1) including both study and sample as random effect (study nested within sample in order to deal with potentially non-independent effect sizes coming from the same manuscript or the same sample of participants (Assink & Wibbelink, 2016; Konstantopoulos, 2011). Publication bias was assessed using the Rank correlation test (Begg & Mazumdar, 1994). This approaches may be less appropriate for mixed-effects meta-analysis which include non-independent data points (Assink & Wibbelink, 2016). However, we provide the results of this tests to be consistent with prior meta-analyses. Heterogeneity was assessed using the Brestlow-day test (Cochran, 1954) and the method proposed by Higgins et al. (termed I-squared; Higgins & Thompson, 2002).

We conducted separate sets of analyses to explore the associations of ELA with two metrics of biological aging: pubertal timing and cellular aging. As described above, data on neural development was not reported in a manner across studies that permitted meta-analysis; instead, these results are systematically reviewed. Within each set of analyses, we began by exploring the association of all adversity types (regardless of dimension) with our two domains of biological aging, then examined whether adversity type was a moderator of these associations. If the moderator analysis was significant, we then ran separate sensitivity analyses to examine associations separately by threat, deprivation, and SES to assess associations of each adversity type with biological aging outcomes.

### Moderator Analyses

In cases where effect sizes showed significant heterogeneity, we tested whether demographic or methodological factors moderated the associations between ELA and biological aging. These factors were based on prior literature, and included sample race/ethnicity (% white), sex composition of the sample (% male), metric of pubertal timing (age of menarche OR measure of secondary sex characteristics), and whether the study controlled for BMI (0/1) or other forms of adversity (0/1). For telomere length and DNAm age we examined the use of blood vs. saliva as a potential moderator in our analyses. If there was no information given by the manuscript regarding a specific moderator then they were marked as missing and not included in the moderator analysis. We tested each moderator separately using the moderator flag in the rma.mv function.

## Results

### Pubertal Timing

The 37 studies included in this meta-analysis produced 86 effect sizes and a total of 63,914 participants. Sample sizes ranged from 25 to 16,202 (Median=480). Of the 86 effect sizes, 20 focused on ELA characterized as threat, 21 focused on deprivation, 33 on SES, and 12 used a cumulative approach of summing across multiple forms of adversity. Table 1 presents descriptive demographic information for each study.

**Table 1.** Characteristics of Individual Studies Included in the Pubertal Timing Meta-Analysis

|  |  |  |  |  |  |  |  |  |
| --- | --- | --- | --- | --- | --- | --- | --- | --- |
| Study | Sample | Sample Size (N) | Age Range | Sex Distribution | Early Life Adversity | Adversity Details | Pubertal Timing Metric | Covariates |
| Amir, Jordan & Bribiescas, 2016 | Nationally Representative Sample (US); Add Health Survey Participants | 3606 | 12-20 | Female only | Threat/SES | Subjective ratings of the availability of resources (family income) and environmental safety | Age at Menarche | None |
| Arim, Shapka, Dahinten, Willms, 2007 | Nationally Representative Sample (Canada); National Longitudinal Survey of Children and Youth | 7,977 | 10-17 | 50% Female | SES | Family SES (composite of household income, parental education and occupation levels, weighted by the factor loadings obtained through principal components factor analysis with varimax rotation) | PDS | Sex |
| Boynton-Jarrett & Harville, 2012 | Nationally Representative Sample (UK); National Child Development Study | 4,524 | 11-16 | Female only | Threat/Deprivation | Exposure to violence/mental health issues; Lack of supportive caregiving | Menarche < 11 years | None |
| Braithwaite, Moore, Lustig, Epel, Ong, Rehkopf, Wang, Miller, & Hiatt, 2009 | Nationally Representative Sample (US); National Growth and Health Study | 2,077 | 9-19 | Female only | SES | SES (parent education level and household income) | Age at Menarche < 12 years | None |
| Colich, Platt, Keyes, Sumner, Allen & McLaughlin | Nationally Representative Sample (US); National Comorbidity Study - Adolescent Sample | 4,937 | 13-18 | Female only | Threat/Deprivation | Threat exposure composite, Deprivation exposure composite | Age at Menarche | None |
| Culpin, Heron, Araya, Melotti, Lewis, & Joinson, 2014 | Nationally Representative Sample (UK); Avon Longitudinal Study of Parents and Children | 3,776 | 8-18 | Female only | SES | Presence of financial problems at age 6 and age 8 and Mother's education status | Age at Menarche | None |
| Deardorff, Abrams, Ekwuru & Rehkopf, 2014 | Nationally Representative Sample (US); National Longitudinal Study of Youth | 4,851 | 9-16 | Female only | SES | SES before age 7 (Family income; Family wealth [income - total debt]; mother's education level; mother's employment status) | Age at Menarche | None |
| Deardorff, Ekwuru, Kushi, Ellis, Greenspan, Mirabedi, Landaverde, & Hiatt, 2011 | Cohort Study of Young Girls' Nutrition, Environment, and Transitions | 444 | 6-10 | Female only | SES | Family Income (<75,000) | Tanner Stage | None |
| Deardorff, Fyfe, Ekwuru, et al 2012 | Cohort Study of Young Girls' Nutrition, Environment, and Transitions | 213 | 6-12 | Female only | SES | Family Income (<50,000) | Tanner Stage | None |
| Ellis & Essex, 2007 | Wisconsin Study of Families and Work | 180 | 0-11 | Female only | SES | SES | PDS and Tanner | None |
| Furu, 1976 | Community sample from Sweden | 917 | 10-17 | Female only | SES | Father's or Mother's Occupation | Age at Menarche | General Health; BMI |
| Graber, Brooks-Gunn & Warren, 1995 | Premenarcheal subset from a larger longitudinal sample | 75 | 10-14 | Female only | General | Number of stressful life events | Age at Menarche and Tanner Stage | None |
| Hartman, Li, Nettle, Belsky, 2017 | Nationally Representative Sample; NICHD Study of Early Child Care and Youth Development (SECCYD) | 659 | 1-15 | Female only | General | Composite of maternal sensitivity, maternal harshness, unpredictability, income harshness | Age at Menarche | General Health; BMI |
| Hayes & Tan, 2016 | Cohort of adopted Chinese girls | 16,202 | 11-13 | Female only | Deprivation | International Adoption (from China to North America) | Age at Menarche | None |
| Johnson, Tang, Almas et al., 2018 | Bucharest Early Intervention Project (BEIP) | 107 | 0-12 | 49% Female | Deprivation | Institutionalization | Age at Menarche and Tanner Stage | None |
| Kelly, Zilanawala, Sacker, Hiatt & Viner, 2017 | Longitudinal cohort from UK Millennium Cohort Study | 5,839 | 0-11 | Female only | SES | SES (Income Quartiles) | Menarche status at age 11 | None |
| Lynne-Landsman, Graber & Andrews, 2010 | Longitudinal cohort from Oregon Youth Substance Use Project; 538 females; 537 males | 1,075 | Grades 1-12 | 52% female | SES | Household resources, conflict and stability | PDS | None |
| Mendle, Leve, Van Ryzin, & Natsuaki, 2014 | Longitudinal intervention cohort of girls in foster care | 100 | 10-12 | Female only | Threat/Deprivation | Sexual Abuse, Neglect, Physical Abuse, Foster Care Placement | PDS | None |
| Mendle, Leve, Van Ryzin, Natsuaki, & Ge, 2011 | Longitudinal intervention cohort of girls in foster care | 100 | 10-12 | Female only | Threat/Deprivation | Sexual Abuse, Neglect, Physical Abuse, Foster Care Placement | Age at Menarche and PDS | All forms of adversity, child's age, child's race |
| Mendle, Moore, Briley & Harden, 2016 | Nationally Representative Sample (US); Add Health Survey Participants | 1,260 | 11-21 | Female only | SES | SES (parental education) | Age at Menarche | None |
| Mendle, Ryan, McKone, 2016 | Nationally Representative Sample (US); Add Health Survey Participants | 6,273 | 11-34 | Female only | Threat/Deprivation/SES | Sexual Abuse, Physical Abuse, Physical Neglect, Undetermined Maltreatment, SES (income to needs ratio) | Age at Menarche and Secondary Sex Characteristics (Breast and Curve Development) | None |
| Milicer, 1968 | Community sample in Wroclaw, Poland | 5,859 | 9-17 | Female only | SES | SES (parental education, occupation and income) | Age at Menarche | age, BMI, race |
| Negriff & Trickett 2012 | Longitudinal cohort in US | 454 | 9-13 | 46.9% female | General | Active case within Department of Child and Family Services | Tanner Stage and PDS | None |
| Negriff, Blankson & Trickett, 2015 | Longitudinal cohort in US | 454 | 9-13 | 47% female | Threat | Sexual Abuse, Other Maltreatment | PDS | ethnicity, obesity, biological father absence |
| Negriff, Saxbe, & Trickett, 2015 | Longitudinal cohort in US | 454 | 9-13 | 47% female | General | Active case within Department of Child and Family Services | PDS | None |
| Noll, Trickett, Long et al., 2017 | Longitudinal cohort in US | 173 | 11-20 | Female only | Threat | Sexual Abuse | Tanner Stage | None |
| Reagan, Salsberry, Fang, Gardner & Pajer, 2012 | Nationally Representative Sample (US); National Longitudinal Study of Youth | 2,337 | 12-17 | Female only | SES | % time in poverty between ages 0-5 | Age at Menarche | All forms of adversity, child's age, child's race |
| Reid, Miller, Dorn, Desjardins, Donzella & Gunnar, 2017 | Cohort of interntionally adopted youth | 165 | 7-14 | Female only | Deprivation | Institutionalizati on | % Menstruating | BMI, people in household, father's occupation |
| Ryan, Mendle & Markowitz, 2015 | Nationally Representative Sample (US); Add Health Survey Participants | 6,273 | 11-34 | Female only | Threat/Depriva tion/SES | Sexual Abuse, Physcial Abuse, Physical Neglect, SES (income to needs ratio) | Age at Menarche | None |
| Salces, Rebato, Susanne, San Martin & Rosique, 2001 | Community sample from Basque Country, Spain | 196 | 9-17 | Female only | SES | SES (father's occupation) | Age at Menarche | None |
| Sheppard, Pearce & Sear, 2016 | Longitudinal sample (UK); Newcastle Thousand Families Study | 251 | 0-15 | Female only | SES | SES (father's occupation) | Age at Menarche | Sex, race/ethnicity, family poverty status, other exposure composite |
| Sonuga-Barke, Schlotz, & Rutter, 2010 | Cohort of adopted Romanian youth | 197 | 6-15 | ~ 50% Female | Deprivation | Intitutionalizati on | Tanner Stage | Parental early puberty history, perinatal factors, childhood and current adiposity, physical activity, sleep duration, presence of stepfather, stressful life events |
| Sumner, Colich, Uddin, Armstrong & McLaughlin, 2018 | Community sample of maltreated children and adolescents from Seattle, WA | 247 | 8-16 | 48% female | Threat/Deprivation/SES | Threat exposure composite, Deprivation exposure composite, SES (below or above poverty line) | Tanner Stage | Sex, Race/Ethnicity, Family Poverty Status, Other Adversity Exposure Composite |
| Sun, Mensah, Azzopardi, Patton & Wake, 2017 | Nationally Representative Sample (Australia); The Longitudinal Study of Australian Children | 3,764 | 0-11 | 48% female | SES | SES (family income, education and occupation) | PDS | None |
| Sung, Simpson, Griskevicius, Kuo, Schlomer & Belsky, 2016 | Nationally Representative Sample (US); NICHD Study of Early Child Care and Youth Development (SECCYD) | 492 | 0-15 | Female only | SES | SES (income to needs ratio) in the first 5 years of life | Age at Menarche | BMI, race, age, other adversity |
| Teilmann, Pedersen, Gormsen, Damgaard, Skakkebaek & Jensen, 2009 | Cohort of Internationally adopted individuals in Denmark | 1,376 | 4-13 | Female only | Deprivation | International Adoption (from Asia, South America, Eastern Europe and Africa to Denmark) | Tanner Stage | None |
| Turner, Runtz & Galambos, 1999 | Canadian cohort of girls recruited from outpatient clinical agencies and public schools | 44 | 12-19 | Female only | Threat | Sexual Abuse | Age at Menarche | None |

#### All Adversities

We first examined the effect of all forms of adversity on pubertal timing across all 37 studies included in the meta-analysis. Greater exposure to ELA was associated with earlier pubertal timing (d=-0.12, 95% CI [-0.22, -0.02]) and significantly differed from zero (Z=-2.41, *p*=0.02; Figure 2). Significant heterogeneity was observed across studies (*Q*(85)=625.50, *p*<0.0001; *I*^2^=96.04). The result of Begg’s publication bias test was not significant (Kendall’s tau=-0.10, *p*=0.18), suggesting no publication bias in our sample of studies.

**Figure 2.**
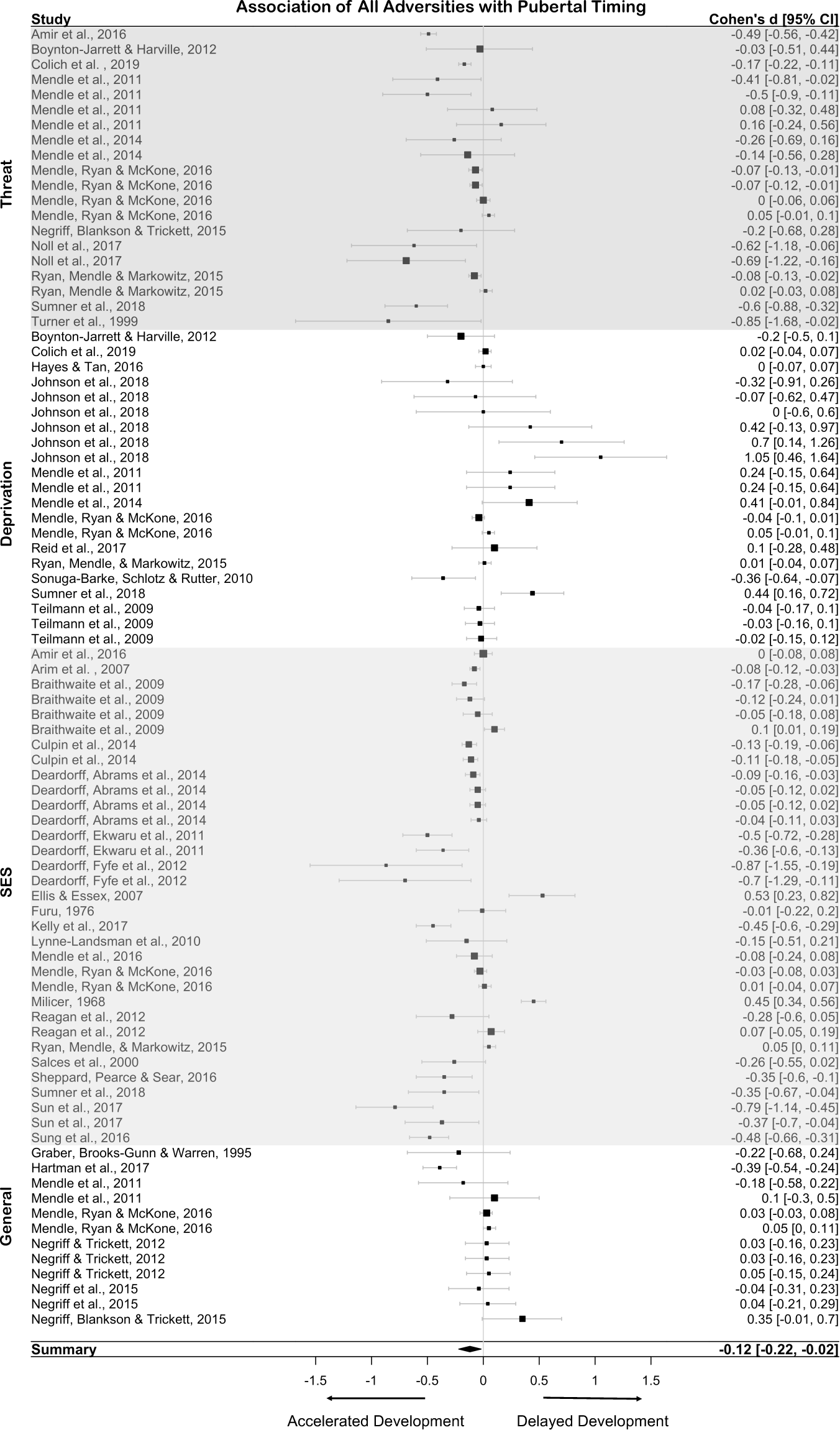
Association of all adversities and pubertal timing.

Using adversity type as a moderator, we tested our hypothesis that threat would have a significant negative effect on pubertal timing (suggesting accelerated development), whereas deprivation would a significant positive effect on pubertal timing (suggesting delayed development). The random-effects meta-analysis including 4 adversity types as a moderator (threat, deprivation, SES, and any studies using only a composite measure of adversity) revealed that adversity type significantly moderated the association between ELA and pubertal timing (*QM*(3)=50.89, *p*<0.0001). Given significant differences across adversity type, we explored the effect of adversity on pubertal timing separately for each category of adversity.

#### Threat

In studies that specifically explored the association of threat exposure with pubertal timing (11 studies; 20 effect sizes, N=20,401), greater exposure to threat was associated with earlier pubertal timing (d=-0.26, 95% CI [-0.41, -0.11]). The effect size was small and significantly differed from zero (Z=-3.44, *p*<0.001; Figure 3). Significant heterogeneity was observed across studies (*Q*(19)=226.32, *p*<0.001; *I*^2^=95.15). The result of Begg’s publication bias test was not significant (Kendall’s tau=-0.31, *p*=0.07), suggesting no publication bias in our sample of studies.

**Figure 3.**
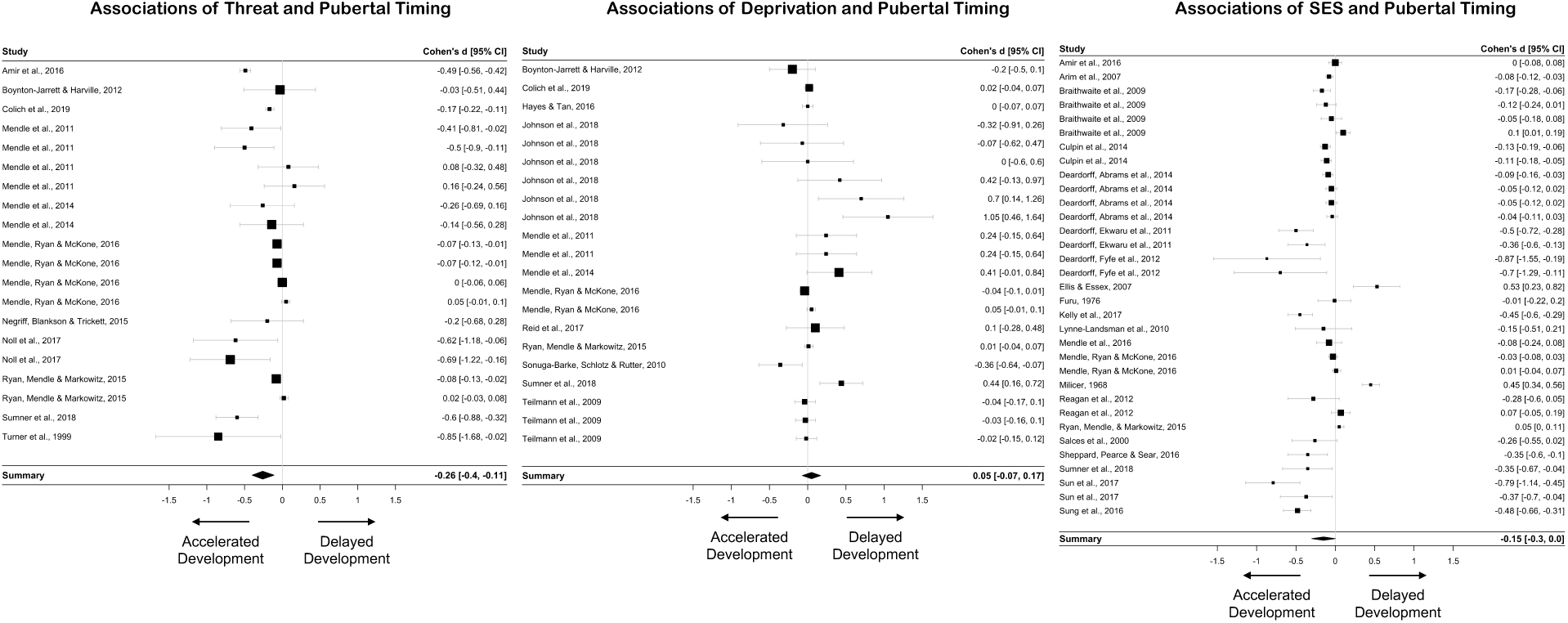
Association of adversity and pubertal timing by adversity type.

Given significant heterogeneity in our studies examining the association of threat-related adversities with pubertal timing, we conducted a series of moderator analyses. None of the five moderators (metric of pubertal timing, sex, race/ethnicity, BMI, controls for other ELA types) were significantly associated with variations in effect size.

#### Deprivation

In studies that specifically focused on the association of deprivation exposure with pubertal timing (12 studies; 21 effect sizes, N=34,193), deprivation was not associated with pubertal timing (d=0.05, 95% CI [-0.07, 0.18]) and did not significantly differ from zero (Z=0.81, *p*=0.42; Figure 3). Significant heterogeneity was observed across studies (*Q*(20)=51.17, *p*<0.001; *I*^2^=89.34). The result of Begg’s publication bias test was not significant (Kendall’s tau= 0.24. *p*=0.15), suggesting no publication bias in our sample of studies.

Given significant heterogeneity in our studies examining the association of deprivation-related adversities with pubertal timing, we conducted moderator analyses. For the association between deprivation and pubertal timing, sex was significantly associated with variation in effect sizes (Estimate=0.01, SE=0.00, Z=3.50, p<0.001), suggesting that the more males included in the sample, the more positive the association between deprivation exposure and pubertal timing (i.e., the more delayed the pattern of maturation).

#### SES

When looking at studies that explored the effect of SES specifically on pubertal timing (21 studies; 33 effect sizes, N=34,489), the random-effects meta-analysis found that SES was associated with earlier pubertal timing (d=-0.15, 95% CI [-0.30, 0.01]), but the effect size was not significantly different than zero (Z=-1.90 *p*=0.06; Figure 3). Significant heterogeneity was observed across studies (*Q*(32)=211.66, *p*<0.001; *I*^2^=96.22). The result of Begg’s publication bias test was significant (Kendall’s tau=-0.27, *p*=0.03), suggesting publication bias in our sample of studies.

Given significant heterogeneity in our studies examining the association of SES with pubertal timing, we conducted a series of moderator analyses. None of the five moderators were significantly associated with variation in effect sizes.

### Cellular Aging

A total of 9 studies (7 examining telomere length, 2 examining DNA methylation age) produced 14 effect sizes across a total of 1,011 participants. Sample sizes for included studies ranged from 38 to 247 (Median=99). Of the 14 effect sizes, 4 focused on the effect of threat on cellular aging, 2 focused on deprivation, 5 on SES, and 3 used a cumulative approach of summing across multiple forms of adversity. Table 2 presents descriptive demographic information for each study.

**Table 2.** Characteristics of Individual Studies Included in the Cellular Aging Meta-Analysis

|  |  |  |  |  |  |  |  |  |
| --- | --- | --- | --- | --- | --- | --- | --- | --- |
| Study | Sample | Sample Size (N) | Age Range | Sex Distribution | Early Life Adversity | Adversity Details | Cellular Aging Metric | Covariates |
| Asok, Bernard, Roth, Rosen & Dozier, 2013 | Longitudinal cohort examining efficacy of attachment-based intervention for infants in Child Welfare System | 38 | 4-7 | 58% Female | General | Identified by Child Welfare System as being at-risk for maltreatment (conditions noted most often include child neglect, domestic violence, parental substance use, inadequate housing) | Telomere Length (buccal swabs) | Household Income, Sex, Minority Status |
| Drury, Mabile, Brett et al. 2014 | Community sample from New Orleans, Louisiana | 80 | 5-15 | 49% Female | Threat/General | Family Instability (witnessing family violence, suicide, incarceration) | Telomere Length (buccal swabs) | None |
| Drury, Theall, Gleason et al., 2012 | Bucharest Early Intervention Project (BEIP) | 100 | 6 months - 10 | 41% Female | Deprivation | Early Institutionalization (% of time spend in institution at 54 months) | Telomere Length (buccal swabs) | Group, sex, ethnicity, age at telomere collection and low birth weight |
| Jovanovic, Vance, Cross et al., 2016 | Longitudinal cohort of trauma exposed children and adolescents | 101 | 6-13 | 54% Female | Threat | Violence Exposure | Horvath DNAm age (saliva) | None |
| Mitchell, Hobcraft, McLanahan, et al., 2014 | 2 cohorts including Fragile Families and Child Wellbeing Study and an independent cohort | 40 | 9 | Male only | General/SES | Harsh Environment determined by equally weighted combination of (i) family economic conditions, (ii) parenting practices, and<br>(iii) family<br>structure/stability -- then<br>incorporating<br>maternal<br>depression;<br>Income to<br>Needs Ratio | Telomere Length (saliva) | None |
| Needham,<br>Fernandez, Lin,<br>Epel &<br>Blackburn, 2012 | Admixture<br>Mapping for<br>Ethnic and<br>Racial Insulin<br>Complex<br>Outcomes<br>(AMERICO)<br>cohort | 70 | 7-13 | 53% Female | SES | SES (family<br>income and<br>parental<br>education) | Telomere<br>Length (blood) | None |
| Shalev, Moffitt,<br>Sugden et al.,<br>2013 | Environmental-Risk<br>Longitudinal<br>Twin Study | 236 | 5-12 | 49% Female | Threat | Domestic<br>Violence,<br>Bullying<br>Victimization,<br>Physical<br>Maltreatment | Telomere<br>Length (buccal<br>swabs) | None |
| Sumner, Colich,<br>Uddin,<br>Armstrong &<br>McLaughlin,<br>2018 | Community<br>sample of<br>maltreated<br>children and<br>adolescents<br>from Seattle,<br>WA | 247 | 8-16 | 48% Female | Threat/Deprivation/SES | Threat exposure<br>composite,<br>Deprivation<br>exposure<br>composite, SES<br>(below or above<br>poverty line) | Horvath DNAm<br>age (saliva) | Sex,<br>Race/Ethnicity,<br>Family Poverty<br>Status, Other<br>Adversity<br>Exposure<br>Composite |
| Theall, Brett,<br>Shirtcliff, Dunn<br>& Drury, 2013 | Community<br>sample from<br>New Orleans,<br>Louisiana | 99 | 4-14 | 53% Female | SES | Household<br>socioeconomic<br>position | Telomere<br>Length (saliva) | None |

**Table 3.** Characteristics of Individual Studies Included in the Cortical Thickness Systematic Review

|  |  |  |  |  |  |  |  |  |  |  |
| --- | --- | --- | --- | --- | --- | --- | --- | --- | --- | --- |
| Authors | Sample | Sample Size (N) | Age Range<sup>1</sup> | Sex Distribution | Early-Life Adversity | Adversity Details | ROI or Whole Brain? | Brain regions with significant associations with adversity) | Covariates | Group Matching Characteristics |
| Busso, McLaughlin, Brueck, Peverill, Gold & Sheridan, 2017 | Community based sample of children in Boston, MA exposed to high rates of maltreatment | 51 | 13-20 | 60% Female | Threat | Childhood Trauma Questionnaire and Childhood Experiences of Care and Abuse to assess emotional, sexual, and physical abuse | ROI (vmPFC, lateral OFC, anterior and posterior cingulate cortices, ventrolateral PFC (including inferior frontal gyrus [IFG]), dorsolateral PFC (including middle and superior frontal gyri), insular cortex, parahippocampal gyrus, temporal pole, and lateral temporal cortex (spanning inferior, middle, and superior temporal gyri) | Thinner cortex among abused in vmPFC, right inferior frontal gyrus, left and right parahippocampal gyri, right inferior temporal gyrus, and right middle temporal gyrus compared to non-exposed controls | age, gender, parental education | race, IQ, handedness |
| Gold, Sheridan, Peverill, Busso, Lambert, Alves, Pine & McLaughlin, 2016 | Community based sample of children in Boston, MA exposed to high rates of maltreatment | 58 | 13-20 | 60% Female | Threat | Childhood Trauma Questionnaire and Childhood Experiences of Care and Abuse to assess emotional, sexual, and physical abuse | ROIs in medial and lateral prefrontal cortex, and medial and lateral temporal cortex | Thinner cortex among abused sample in bilateral ventromedial prefrontal cortex, right orbitofrontal cortex, left temporal pole, right inferior frontal gyrus, left and right parahippocampal gyri, left and right inferior temporal gyrus, right superior temporal gyrus, and right middle temporal gyrus | race, age, sex, and parental education |  |
| McLaughlin, Sheridan, Gold, Duys, Lambert, Peverill, Heleniak, Shechner, Wojcieszak | Community based sample of children in Seattle, WA exposed to high rates of | 60 | 18-Jun | 48% Female | Threat | Childhood Trauma Questionnaire and Childhood Experiences of Care and Abuse to assess | ROI in dorsal anterior cingulate and ventromedial prefrontal cortex | No significant associations with cortical thickness | age, sex |  |
| & Pine, 2016 | maltreatment |  |  |  |  | emotional, sexual, and physical abuse |  |  |  |  |
| Kelly, Viding, Wallace, Schaer, De Brito, Robustelli & McCrory, 2013 | Community based sample recruited from London Social Services department | 43 | 12.77(1.19) for controls, 12.27(1.41) for maltreated | 56% Female | Threat | Social worker report using Kaufman's 4 point scale for physical abuse, emotional abuse, sexual abuse, and neglect | Whole brain | decreased cortical thickness in mPFC in maltreated, | age and sex | Tanner stage, handedness, cognitive ability, ethnicity |
| Lawson, Duda, Avants, Wu & Farah, 2013 | Participants from NIH Pediatric MRI Data Repository | 283 | 11.47 (3.50) | 53% Female | SES | Family income, parental education summed from two parents. Parental education variable was square-root transformed | ROIs in the prefrontal cortex | Lower parental education was associated with decreased cortical thickness in right anterior cingulate cortex and left superior frontal gyrus. No associations emerged with parental income. | age, sex, total brain volume, race, body mass index | N/A |
| Mackey, Finn, Leonard, Jacoby-Senghor, West, | Community sample from Boston, Massachusetts area | 58 | Lower income: 14.47 (0.38), Higher income: | 53% Female | SES | Free or reduced-price lunch status obtained from the | Whole brain | Thinner cortex in lower income group in bilateral | sex, intracranial volume. Associations also hold when | None |
| Gabrieli, Gabrieli, 2015 |  |  | 14.35 (0.47) |  |  | Massachusetts Department of Elementary and Secondary Education |  | temporal and occipital lobes and in right lateral prefrontal cortex | controlling for race and ethnicity |  |
| Jednoróg, Altarelli, Monzalvo, Fluss, Dubois, Billard, Dehaene-Lambertz & Ramus, 2012 | Community sample from Paris, France | 23 | 8-10 | 57% Female | SES | Hollingshead Two-Factor Index of Social Position using maternal report of maternal occupation and education | Whole Brain | Positive associations between SES and thickness in L angular gyrus, L heschl's gyrus, L post-ventral cingulate, and L anterior occipital sulcus. Negative associations between SES and thickness in L fronto-maginal gyrus/sulcus, L middle frontal sulcus, and L intraparietal sulcus. | None | N/A |
| Noble, Houston, Brito, Bartsch, Kan, Kuperman, Akshoomoff, Amaral, Bloss, Libiger, Schork, Murray, Casey, Chang, Ernst, Frazier, Gruen, Kennedy, Van Zijl, Mostofsky, Kaufmann, Kenet, Dale, Jernigan & Sowell, 2015 | Pediatric Imaging, Neurocognition, Genetics Study (PING) | 1148 | 3-20 | 48% Female | SES | parent report of income and education | Whole Brain | No significant associations between cortical thickness and SES or SES x Age interactions | age, age-squared, scanner, sex, genetic ancestry | N/A |
| Piccolo, Merz, He, Sowell, Noble KG; Pediatric Imaging, Neurocognition, Genetics Study, 2016 | Pediatric Imaging, Neurocognition, Genetics Study (PING) | 1148 | 3-20 | 48% Female | SES | parent report of income and education | Whole Brain | Age(square d) X SES (both income and education) interaction in average cortical thickness, such that lower SES children demonstrate faster thinning across the whole cortex. No specific regions survive correction for multiple comparisons | SES measure, age, age-squared, scanner, sex, genetic ancestry, and parental SES measure x age | N/A |
| Rosen, Sheridan, Sambrook, Meltzoff & McLaughlin, 2018 | Community based sample of children in Seattle, WA exposed to high rates of maltreatment (†) | 53 | 6-19 | 48% Female | Deprivation/ SES | SES-Family income divided by the federal poverty line for a family of that size; Cognitive stimulation-Parent report using Home Observation Measurement of the Environment (HOME)-short form | Whole Brain and ROI | Lower levels of cognitive stimulation were associated with thinner cortex in left superior parietal lobule and left middle frontal gyrus. There were no associations with SES and no significant whole brain associations. | age, sex, violence exposure | N/A |
| Avants, Hackman, Betancourt, Lawson, Hurt & Farah, 2015 | 52 young adults | 52 | 19.16 years (SD = 1.32) | 52% female | Deprivation | Environmental Stimulation as measured by the HOME at ages 4 and 8 | Whole Brain | Thicker cortex among children with lower levels of cognitive stimulation at age 4 (but not 8) in L lateral inferior temporal—posterior, | age at scan, maternal IQ, gender, foster care and prenatal cocaine | N/A |
|  |  |  |  |  |  |  |  | L lateral<br>Inferior<br>temporal<br>-anterior,<br>Bilateral<br>fusiform, R<br>lateral<br>inferior<br>temporal |  |  |
| Hodel,<br>Hunt,<br>Cowell, Van<br>Den<br>Heuvel,<br>Gunnar &<br>Thomas,<br>2015 | 110 post-<br>institutionali<br>zed<br>children,<br>from<br>Minnesota<br>Internationa<br>l Adoption<br>Project | 172 | 12-14 | 61% female | Deprivation | previously<br>institutionali<br>zed<br>children<br>adopted<br>into the<br>United<br>States | Whole<br>Brain | Thinner<br>cortex in<br>right inferior<br>frontal<br>gyrus<br>among<br>previously<br>institutionali<br>zed<br>children<br>compared<br>to controls | age, sex |  |
| McLaughlin<br>, Sheridan,<br>Winter,<br>Fox,<br>Zeanah &<br>Nelson,<br>2014 | Bucharest<br>Early<br>Intervention<br>Project<br>(BEIP) | 80 | 8-10 | 50% female | Deprivation | part of a<br>randomized<br>control trial<br>where 1/2<br>of the<br>institutionali<br>zed<br>children<br>were<br>placed in<br>high quality<br>forster care.<br>There were<br>no<br>differences<br>between<br>the care as<br>usual group<br>and foster<br>care group,<br>so all<br>analyses<br>compared<br>the Ever | Whole<br>Brain | widespread<br>reductions<br>in cortical<br>thickness<br>across<br>frontal,<br>parietal and<br>temporal<br>lobes in<br>children<br>who were<br>ever<br>institutionali<br>zed | age, sex,<br>total brain<br>volume,<br>and TR<br>length (bc<br>different<br>acquisition<br>parameters<br>) |  |
|  |  |  |  |  |  | Institutionalized Group (EIG) to the Never Institutionalized Group (NIG) |  |  |  |  |
| <sup>1</sup>Age range (in. years) was the most commonly used descriptor of the full sample. When this was not available, means and standard deviations are given. |  |  |  |  |  |  |  |  |  |  |

**Table 4.** Characteristics of Individual Studies Included in the amygdala-mPFC Review

|  |  |  |  |  |  |  |  |  |  |  |  |
| --- | --- | --- | --- | --- | --- | --- | --- | --- | --- | --- | --- |
| Authors | Sample | Sample Size (N) | Age Range | Sex Distribution | Early-Life Adversity | Adversity Details | Seed-to-seed or Whole Brain? | Task | Associations with adversity and amygdala-mPFC connectivity | Covariates | Group Matching Characteristics |
| Cisler, Scott, Smitherman, Lenow & Kilts, 2013 | Community based sample in Arkansas | 30 | 12-16 | 100% female | Threat | Assessed using the trauma assessment section of the National Survey of Adolescents (sexual assault, physical assault, severe abuse from caregiver) | whole brain | implicit emotion processing task | greater connectivity between right amygdala and left middle frontal gyrus, and right amygdala and left perigenual anteriorcingulate | none | Age, ethnicity, anxiety and depression diagnoses |
| Colich, Williams, Ho et al., 2017 | Community based sample in the San Francisco Bay Area | 137 | 9-13 | 57% female | Threat | Based on modified version of Traumatic Events Screening Inventory for Children | seed-to-seed | implicit emotion regulation task | ELS was associated with increased negative connectivity between the right ventrolateral prefrontal cortex and bilateral amygdala | age | N/A |
| Gee, Gabard-Durnam, Flannery et al., 2013 | Post institutionalized children adopted into the United States and comparison children* | 89 | 6-18 | 56% female | Deprivation | Sample of previously institutionalized children adopted into the United States | whole brain | Emotional face viewing task | Comparison group children showed positive amygdala-mPFC connectivity during viewing of fearful faces while adolescents showed comparison group adolescents showed negative connectivity. In contrasts, both PI children and adolescents showed negative amygdala-mPFC connectivity | amygdala reactivity | Pubertal status |
| Keding & Herringa, 2016 | Community based sample in Wisconsin. Sampled specifically for PTSD | 53 | 8-18 | 58% female | Threat | All trauma exposed participants also had PTSD. Trauma exposures were Sexual abuse, traumatic death of loved one, witnessing violence, accident, physical abuse | whole brain | implicit dynamic emotion face | PTSD sample showed reduced connectivity between amygdala and both dorsal anterior cingulate and ventrolateral PFC during angry trials and increases connectivity to happy faces. Control participants showed the opposite pattern of results. | age, sex | age, pubertal status, IQ, handedness |
| Marusak, Martin, Etkin & Thomason, 2015 | Community based sample in Detroit, MI | 30 | 9-17 | 80% female | Threat | At least one traumatic life event endorsed on parent or child report of Children's Trauma Assessment Center Screen Checklist. These included sexual abuse, physical abuse, neglectful home environment, exposure to domestic violence, exposure to other violence, multiple separations from caregiver | whole brain | Emotional Stroop | During emotional conflict regulation, trauma exposed group demonstrated positive amygdala - pregenual cingulate cortex connectivity while the control group demonstrated negative connectivity. | none | age, sex, IQ |
| Silvers,<br>Lumian,<br>Gabard-Durnam et<br>al., 2016 | Post<br>institutionaliz<br>ed children<br>adopted into<br>the United<br>States and<br>comparison<br>children* | 89 | 7-16 | 62% female | Deprivatio<br>n | Sample of<br>previously<br>institutionaliz<br>ed children<br>adopted into<br>the United<br>States | whole brain | Aversive<br>learning<br>paradigm | During a<br>fear<br>conditioning<br>task, PI<br>youth<br>demonstrate<br>positive<br>connectivity<br>between the<br>amygdala<br>and vmPFC,<br>dmPFC, and<br>vIPFC ,<br>while the<br>compairosin<br>group<br>demonstrate<br>s negative<br>connectivity | average<br>head<br>motion, | age, IQ |

#### All Adversities

We first examined the effect of all forms of adversity on cellular aging across all 9 studies included in the meta-analysis. The random-effects meta-analysis found that greater exposure to early-life adversity was associated with accelerated cellular aging (d=-0.30, 95% CI [-0.46, -0.15]) and significantly differed from zero (Z=-3.83, *p*<0.0001; Figure 4). There was no significant heterogeneity observed across studies (*Q*(13)=20.30, *p*=0.09; *I*^2^=31.64. The result of Begg’s publication bias test was not significant (Kendall’s tau=-0.27, *p*=0.19), suggesting no publication bias in our sample of studies.

**Figure 4.**
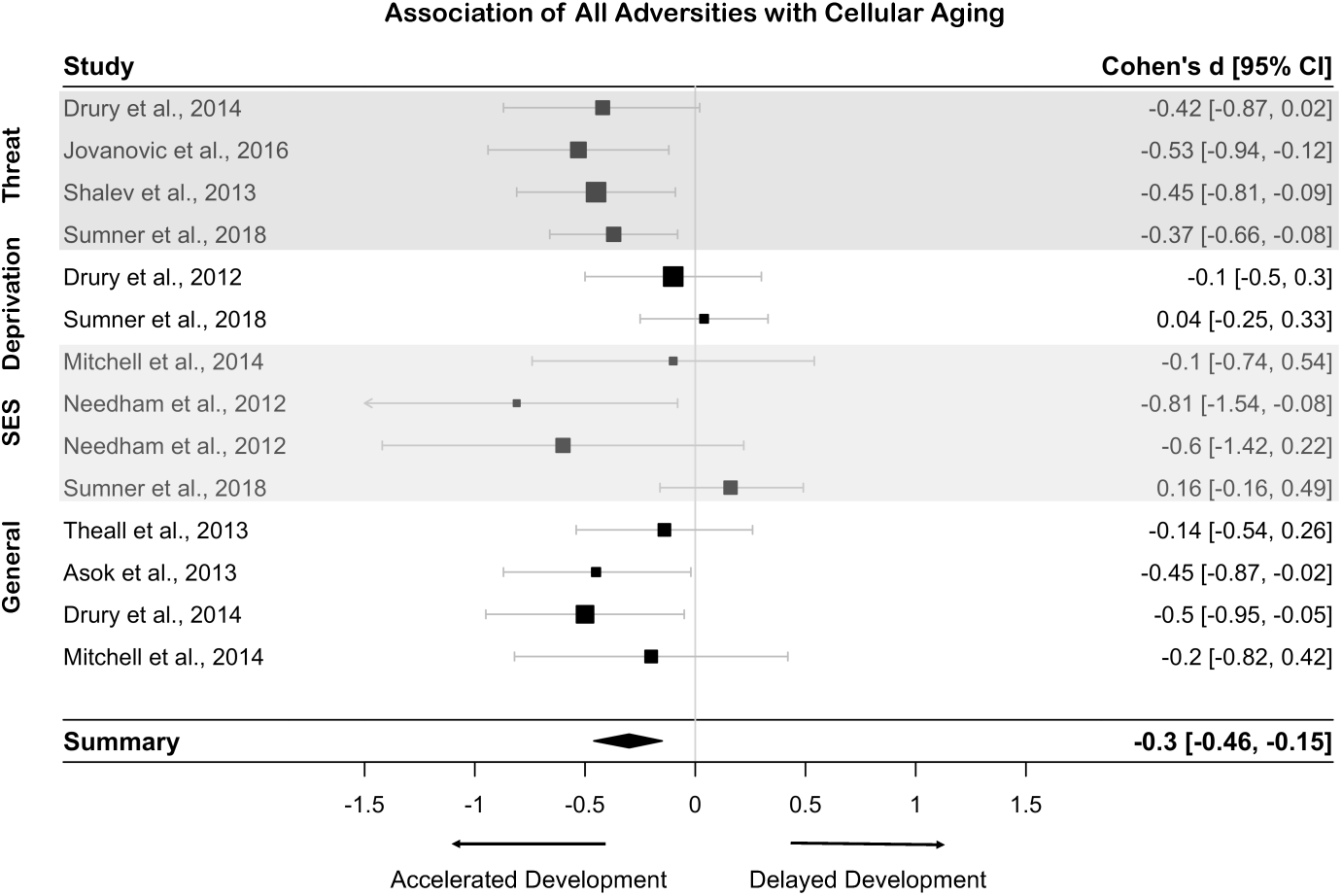
Association of all adversities and cellular aging.

Although there was limited heterogeneity across studies, given our theoretical interests and hypotheses about differences in associations by adversity type, we examined adversity type as a moderator of these associations to evaluate our hypothesis that threat would be associated with advanced cellular aging and determine whether these effects were similar for other adversity types. Adversity type significantly moderated the association between ELA and cellular aging (*QM*(3)=9.68, *p*=0.02). We additionally explored the associations of ELA with cellular aging separately for each adversity type.

#### Threat

In studies that explored the association of threat exposure with cellular aging (4 studies; 4 effect sizes, N=664), greater exposure to threat was associated with accelerated cellular aging (d=-0.43, 95% CI [-0.61, -0.25]). The effect size was moderate in magnitude and differed from zero (Z=-4.65, *p*<0.0001; Figure 5). Significant heterogeneity was not observed across studies (*Q*(3)=0.39, *p*=0.94; *I*^2^=0.00000001). The result of Begg’s publication bias test was not significant (Kendall’s tau=-0.33, *p*=0.75), suggesting no publication bias in our sample of studies. We did not explore moderators given the lack of heterogeneity in effect sizes.

**Figure 5.**
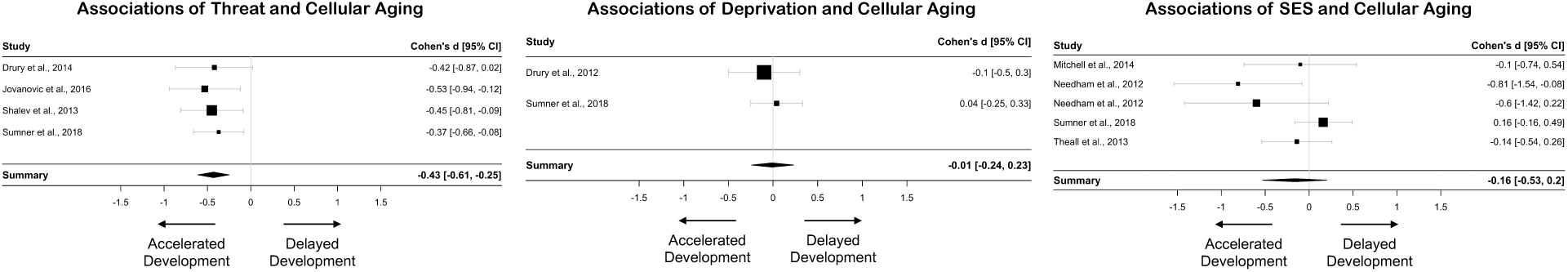
Association of adversity and cellular aging by adversity type.

#### Deprivation

In studies that explored the association of deprivation exposure with cellular aging (2 studies; 2 effect sizes, N=347), the random-effects meta-analysis found that deprivation was not associated with cellular aging (d=-0.06, 95% CI [-0.30, 0.17]), with an effect size that did not significantly differ from zero (Z=-0.51 *p*=0.61; Figure 5). Significant heterogeneity was not observed across studies (*Q*(1)=0.060, *p*=0.81; *I*^2^=0.00000001). The result of Begg’s publication bias test was not significant (Kendall’s tau=-1.000, *p*=1.000), suggesting no publication bias in our sample of studies. However, given this analysis only contained two effect sizes, this is not a reliable estimate of publication bias. We did not explore moderators given the lack of heterogeneity in effect sizes.

#### SES

In studies of SES and cellular aging (4 studies; 5 effect sizes, N=456), SES was not associated with cellular aging (d=-0.16, 95% CI [-0.53, 0.20]), with an effect size that did not significantly differ from zero (Z=-0.87, *p*=0.38; Figure 5). Significant heterogeneity was not observed across studies (*Q*(4)=7.64, *p*=0.11; *I*^2^=53.89). The result of Begg’s publication bias test was not significant (Kendall’s tau=-0.60, *p*=0.23), suggesting no publication bias in our sample of studies. We did not explore moderators given the lack of heterogeneity in effect sizes.

### Brain Development

We systematically reviewed the associations between ELA and two metrics of brain development: cortical thickness and task-based amygdala-PFC functional connectivity. Across the two metrics there were 19 studies across a total of 2,276 unique participants (13 cortical thickness papers, N = 1,848, 6 amgydala-PFC connectivity papers, N = 428).

### Cortical Thickness

#### Threat

We found four papers that investigated the association between experiences of threat and cortical thickness in childhood and adolescence (N=161; Busso et al., 2017; Gold et al., 2016; Kelly et al., 2013; McLaughlin, Sheridan, et al., 2016). Of these four studies, three found that children exposed to threat had accelerated thinning of the cortex. Critically, all three of these studies found decreased cortical thickness among children exposed to threat in the ventromedial PFC (Busso et al., 2017; Gold et al., 2016; Kelly et al., 2013), and two found additional associations that follow the same pattern of decreased thickness among threat-exposed youths in regions including the lateral PFC and medial and lateral temporal cortex (Busso et al., 2017; Gold et al., 2016). In contrast, one study that spanned a larger age range found no association between experiences of threat and cortical thickness in regions of interest in the dorsal anterior cingulate or ventromedial PFC (McLaughlin, Sheridan, et al., 2016). As a whole, these studies provide support for the hypothesis that experiences of threat are associated with accelerated development, especially in the ventromedial PFC.

#### Deprivation

A total of four studies (N = 353) met inclusion criteria for our review of the association between experiences of deprivation and cortical thickness. This included two studies investigating cortical structure among previously institutionalized children (Hodel et al., 2015; McLaughlin, Sheridan, Winter, et al., 2014) and two investigating the association between cognitive stimulation/deprivation in the home environment and cortical structure (Avants et al., 2015; Rosen et al., 2018). Both studies of institutionalized children demonstrate support for the hypothesis that experiences of deprivation are associated with accelerated cortical thinning. In one study institutionalization was associated with widespread reductions in cortical thickness, including in nodes of the frontoparietal and dorsal attention networks (superior parietal lobule, frontal pole, superior frontal gyrus), default mode network (inferior parietal cortex, precuneus, mid-cingulate), lateral temporal cortex, parahippocampal cortex, and insula at age 8-10 years (McLaughlin, Sheridan, Winter, et al., 2014). In contrast, Hodel and colleagues (2015) found reduced cortical thickness only in the inferior frontal gyrus among previously institutionalized children compared to controls at age 12-14 years.

The other two studies investigated cortical thickness and its association with cognitive stimulation/deprivation in the home environment. In a cross-sectional study spanning children and adolescents, cognitive stimulation (i.e., lower deprivation) was positively associated with cortical thickness in two nodes in the left, but not right frontoparietal network (superior parietal lobule and middle frontal gyrus) (Rosen et al., 2018). These results are consistent with the idea that deprivation (i.e. low cognitive stimulation) is associated with accelerated development. In contrast, in a longitudinal study, results revealed that cognitive stimulation at age 4, but not at age 8, was negatively associated with cortical thickness at age 19, such that lower deprivation was associated with thicker cortex in the ventral temporal cortex and inferior frontal gyrus. These results suggest that cognitive deprivation is associated with delayed development in these regions.

#### SES

Six studies met the criteria for inclusion examining SES and cortical thickness (N = 1512) (Jednoróg et al., 2012; Lawson, Duda, Avants, Wu, & Farah, 2013; Mackey et al., 2015; Noble et al., 2015; Piccolo, Merz, He, Sowell, & Noble, 2016; Rosen et al., 2018). Of those, four found that low SES was associated with thinner cortex across large swaths of cortex encompassing the frontoparietal network (lateral prefrontal cortex, superior parietal cortex), default mode network (lateral temporal cortex, precuneus), and the visual system (lateral occipital and ventral temporal cortex), supporting the idea that low SES is associated with accelerated cortical development. Piccolo and colleagues (2016) found that SES moderates the association between age squared and cortical thickness such that low SES individuals show a sharper decline in cortical thickness early in development, which may reflect accelerated development compared to higher SES individuals. Additionally, Lawson and colleagues (2013) demonstrate that low parent education is associated with reduced cortical thickness in the right cingulate gyrus and right superior frontal gyrus, and Mackey and colleagues (2015) demonstrate that low SES individuals demonstrate thinner cortex across much of the brain including the frontoparietal network (right middle frontal gyrus, left superior parietal lobule, right frontal pole), default mode network (left precuneus, bilateral lateral temporal cortex, right frontal pole), and visual system (bilateral occipital and ventral temporal cortex). Two studies spanning larger age ranges (Noble et al., 2015; Rosen et al., 2018) found no association between SES and cortical thickness. Importantly, Noble et al., (2015) and Piccolo et al., (2016) used the same sample and while there were no main effects of SES on cortical thickness and no age x SES interactions (Noble et al., 2015), Piccolo and colleagues demonstrate an age squared x SES interaction such that children from low-income households demonstrate accelerated thinning compared to high-income counterparts.

### Amygdala-PFC Connectivity

#### Threat

Our search yielded four papers that evaluated the association between amygdala-PFC connectivity and threat-related experiences. Of these four studies (N = 250), two support the hypothesis that experiences of threat are associated with accelerated maturation of this network such that threat-exposed children exhibit more negative connectivity between amygdala and PFC during both an implicit dynamic emotion face task, and an explicit affect labeling task, than children of the same age (Colich et al., 2017; Keding & Herringa, 2016). The two other studies demonstrate the opposite pattern of results such that children who have experienced threat demonstrate more positive task-related amygdala-PFC connectivity compared to controls while viewing emotional faces and while performing an emotional conflict task (Cisler, Scott Steele, Smitherman, Lenow, & Kilts, 2013; Marusak et al., 2015). These mixed findings do not provide conclusive evidence that experiences of threat are associated with either accelerated or delayed development of the circuits.

#### Deprivation

Our search yielded two papers (N = 89) that evaluated the association between experiences of deprivation and task-related connectivity between mPFC and amygdala. Of these two studies, one demonstrated evidence for accelerated development of these circuits such that children who have experienced deprivation exhibit more negative connectivity earlier in development than comparison children in a passive viewing task of facial emotion (Gee, Gabard-Durnam, et al., 2013). The other study found the opposite pattern of results such that children who had experienced deprivation demonstrated more positive amygdala-mPFC connectivity than comparison children in a fear conditioning paradigm (Silvers, Lumian, et al., 2016).

## Discussion

Through the use of meta-analysis and systematic review, we provide evidence that ELA accelerates biological aging, as measured by pubertal timing, cellular aging, and cortical thinning in childhood and adolescence. We found no evidence for a consistent effect of ELA on accelerated development of amygdala-mPFC connectivity. First, although we observed an overall association of ELA with pubertal timing, moderator analysis revealed that ELA characterized by threat, but not deprivation or SES, was associated with accelerated pubertal development with a small effect size, suggesting specificity in the link between ELA and pubertal timing to threat-related adversity. Second, ELA was also associated with accelerated cellular aging as measured by both leukocyte telomere length and DNA methylation age. Again, moderator analyses revealed accelerated aging among children exposed to threat of moderate effect size, but no association with deprivation or SES. Finally, the results of our systematic review of the effects of ELA on brain development revealed a consistent association between ELA and accelerated cortical thinning across multiple types of ELA, although the specific brain regions involved vary by adversity type. Associations of threat with cortical thinning were most consistent in ventromedial PFC, whereas associations of deprivation with cortical thinning were most consistent in the frontoparietal and default mode networks and the ventral visual stream. In contrast, there was no consistent association of ELA with amygdala-mPFC connectivity. These findings suggest both common and specific effects of dimensions of ELA across multiple domains of biological aging.

### ELA and Pubertal Timing

ELA was associated with accelerated pubertal timing overall, but significant heterogeneity existed in this effect as a function of adversity type. The strength of the association of ELA with pubertal timing was significantly moderated by adversity type, such that the association between ELA and accelerated pubertal timing was specific to experiences characterized by threat, and showed no association with deprivation or SES. These results are consistent with predictions from life history models that exposure to environmental harshness (i.e. threat) in childhood accelerates sexual maturation, in order to increase chances of reproduction prior to mortality (J. Belsky et al., 2012; Ellis et al., 2009). They are also consistent with recent findings from our lab demonstrating that threat-related adversities are associated with accelerated pubertal development even after adjustment for exposure to co-occurring deprivation (Colich et al., 2019; Sumner et al., 2019). Some have argued that ELA impacts pubertal timing through influences on the hypothalamic-pituitary-adrenal (HPA) axis (Negriff, Saxbe, et al., 2015; Saxbe, Negriff, Susman, & Trickett, 2015). Given associations between threat-related adversity and altered diurnal patterns of cortisol and cortisol reactivity in childhood (Carpenter, Shattuck, Tyrka, Geracioti, & Price, 2011; Jaffee et al., 2014; King et al., 2017; Tyrka et al., 2009), it is plausible that trauma-related alterations of the HPA-axis may interact with the HPG-axis to accelerate the onset of pubertal development (Negriff, Saxbe, et al., 2015; Saxbe et al., 2015). It is also important to consider the role of gene-environment correlation in the association between threat-related ELA and pubertal timing (Cousminer, Widén, & Palmert, 2015; Harden, 2014; Rowe, 2002). For instance, mothers who experience earlier onset of puberty may reproduce at an earlier age, and have children who are both more likely to experience trauma and an earlier onset of puberty (de Vries, Kauschansky, Shohat, & Phillip, 2004; Towne et al., 2005). Future research should explore how maternal age at menarche influences the associations of ELA and pubertal timing in their offspring in order to better understand the mechanisms linking threat-related ELA and accelerated pubertal development.

We did not find support for our hypothesis that ELA characterized by deprivation would show an association with delayed pubertal timing. Instead, we found no association between deprivation and pubertal timing. Life history theory posits that deprivation of bioenergetics resources could result in delayed maturation and later pubertal development (J. Belsky et al., 2012; Ellis et al., 2009). In this analysis, we included emotional and physical neglect (Boynton-Jarrett & Harville, 2012; Colich et al., 2019; Mendle et al., 2014, 2011, 2016; Ryan et al., 2015; Sumner et al., 2019), and early institutionalization (G. et al., 2009; Hayes & Tan, 2016; Johnson et al., 2018; Reid et al., 2017; Sonuga-Barke, Schlotz, & Rutter, 2010) as forms of deprivation. It is likely that deprivation in our modern context, represented by the forms of psychosocial deprivation included in our analyses, is qualitatively different from deprivation in our evolutionary past. Whereas there is strong evidence for associations of food insecurity and severe deprivation associated with war and famine with delayed pubertal timing (Prebeg & Bralic, 2000; van Noord & Kaaks, 1991), there is less support for association of early institutionalization (where most children experience severe emotional deprivation but not necessarily food insecurity) with pubertal timing (Johnson et al., 2018; Reid et al., 2017).

In this meta-analysis, we decided to isolate the effects of deprivation (including neglect and early life institutionalization) from the effects of SES. Low SES, defined as poverty and low parental education, has previously been used as an indicator of deprivation in studies that adjust for co-occurring threat exposure (e.g., Lambert et al., 2017; Sheridan et al., 2017), based on extensive evidence demonstrating that children from families with low parental education and/or income experience reductions in cognitive and social stimulation than children from higher-SES families (Bradley et al., 2001; Duncan & Magnuson, 2012). However, within the DMAP model, poverty is conceptualized as a risk factor for both threat and deprivation, rather than a direct marker of deprivation (McLaughlin, Sheridan, Lambert, et al., 2014; Sheridan & McLaughlin, 2014). Indeed, there is a strong association between SES and exposure to violence (Foster, Brooks-Gunn, & Martin, 2007) in addition to deprivation. Here, the association of SES with accelerated pubertal timing was not significant, potentially reflecting the fact that SES is a risk marker for exposure to other forms of adversity (e.g., trauma) that are associated with accelerated pubertal development rather than having a direct effect on pubertal timing. Overall, these findings highlight the importance of considering the nature of the exposure when exploring the developmental consequences of ELA. Future research should carefully distinguish between the effects of threat- and deprivation-related adversities on pubertal timing.

There was no evidence for the moderating effect of metric of pubertal timing (age of menarche vs. secondary sex characteristics) or race/ethnicity on associations of ELA with pubertal timing, or whether studies controlled for BMI or exposure to other adversity types. However, sex did moderate the association between deprivation-related adversities and pubertal timing such that the more males included in the sample, the more positive the association between deprivation exposure and pubertal timing, suggesting more delayed pubertal maturation. These results suggest that deprivation may have a differential effect on males and females. Animal models of nutritional challenge (including undernutrition and obesity) have differential effects on pubertal timing in male and female rats (Sánchez-Garrido et al., 2013), suggesting that sex-specific metabolic effects of deprivation may have a significant impact on pubertal timing. Future research in humans studying the impact of food insecurity specifically, should explore this question directly.

### ELA and Cellular Aging

ELA was associated with accelerated cellular aging, as measured by both leukocyte telomere length and DNAm age, such that greater exposure to adversity was associated with decreased telomere length and more advanced DNAm age relative to chronological age. These results replicate earlier meta-analyses conducted in adult populations of adversity with DNAm age (Wolf et al., 2018) and telomere length (Hanssen, Schutte, Malouff, & Epel, 2017). These results are also broadly consistent with an earlier meta-analysis exploring the effects of stress exposure (broadly defined) on telomere length (Coimbra et al., 2017). The consistency in findings is striking given significant differences in the approach of these meta-analyses. Whereas Wolf and colleagues (2018) and Hanssen and colleagues (2017) examined the association between ELA and accelerated biological aging in adults, Coimbra et al. (2017) examined a broad range of stressors in childhood and adolescence, including stress reactivity as indexed by cortisol reactivity and parental psychopathology. We did not include cortisol reactivity or parental psychopathology as adversities in the current meta-analysis, yet results are largely consistent with Coimbra and colleagues.

We did not observe heterogeneity in the associations of ELA with cellular aging. However, in stratified analysis, we found that exposure to threat was associated with accelerated cellular aging of moderate magnitude, whereas neither deprivation nor SES was associated with cellular aging. These differential associations should be interpreted with caution, however, as our analysis of whether type of ELA was a moderator effect size magnitude was only significant at a trend-level and the number of studies examining deprivation and SES with cellular aging were small. Nonetheless, these findings suggest that threat-related adversities are consistently associated with accelerated cellular aging. Greater work is needed to clarify the magnitude and direction of effects for deprivation and childhood SES.

These results are consistent with “internal prediction” models of predictive adaptive response (Nettle et al., 2013; Rickard et al., 2014), which propose that ELA negatively influences physical health through altered cellular development as a result of reduced energy to build or repair cellular tissue. This theory expands earlier models focused on allostatic load, or the accumulation of environmental insults on biological systems (Danese & McEwen, 2012; McEwen, 1998; McEwen & Stellar, 1993), and developmental origins of health and disease models (Barker, 2007) focused on how early experience programs biological development to adapt to later environmental conditions. These theories all suggest that accelerated cellular aging occurs as a result of environmental experiences in development. Accelerated cellular aging following ELA may occur in response to alterations in mitochondrial function, oxidative stress, and inflammation (Shalev, 2012). Although we observed consistent effects of ELA across two metrics of cellular aging (telomere length and DNAm age), some work indicates that exposure to ELA may not have consistent associations across other metrics of biological aging (D. W. Belsky et al., 2015). Future research should explore the effect of distinct forms of ELA on additional metrics of allostatic load that may represent accelerated biological aging, including cardiometabolic risk, inflammation, and respiratory health.

### ELA and Brain Development

#### Cortical Thickness

Consistent with the hypothesis that ELA leads to accelerated development, the majority of studies investigating the association between ELA and cortical thickness found that children exposed to adversity of any kind have thinner cortex than their non-exposed counterparts across threat, deprivation, and SES. However, it is critical to note that the specific brain regions that exhibited this pattern of thinning varied consistently by adversity type. This specificity may reflect precocious maturation of particular regions of the brain depending on the particular type of adversity experienced, reflecting adaptive experience-related tuning of neural systems to the environment in which they are developing. There was remarkable consistency across studies of threat-related experiences and cortical thickness, with the majority observing thinner cortex in the ventromedial PFC among children exposed to trauma (Busso et al., 2017; Gold et al., 2016; Kelly et al., 2013). The vmPFC is implicated in multiple forms of emotion processing, including recall of extinction learning, appraisal of episodic memories, and appraisal of simulated future events (Dixon, Thiruchselvam, Todd, & Christoff, 2017; Milad & Quirk, 2012; Phelps & LeDoux, 2005). The vmPFC has strong interconnections with the amygdala and modulates amygdala activation based on appraisals and prior learning (Phelps & LeDoux, 2005). Accelerated thinning of this region among children exposed to trauma could reflect earlier or more frequent recruitment of this region to modulate amygdala responses, which are well-established to be elevated in response to threat cues among children exposed to violence (Hein & Monk, 2017; McCrory, De Brito, & Viding, 2011; McLaughlin, Peverill, Gold, Alves, & Sheridan, 2015), ultimately producing more rapid specialization of this region, potentially through more rapid synaptic pruning or increased myelination in this region.

Association between experiences of deprivation and cortical structure were more mixed. While one study of previously institutionalized children demonstrated widespread reductions in cortical thickness across regions of the frontoparietal, default mode, and visual networks (McLaughlin, Sheridan, Winter, et al., 2014), another found reduced cortical thickness only in the inferior frontal gyrus (Hodel et al., 2015). Studies investigating low cognitive stimulation have also been mixed. While one study found that low cognitive stimulation was associated with thinner cortex in the frontoparietal network across childhood and adolescence (Rosen et al., 2018), another found that lower cognitive stimulation was associated with thicker cortex in the lateral prefrontal cortex and ventral visual stream in late adolescents (Avants et al., 2015). Differences in the age of the samples and timing of assessment of cognitive stimulation may have contributed to these inconsistent findings.

The studies investigating SES-related differences in cortical thickness also had mixed results. Two studies found widespread positive associations with SES and thickness in the frontoparietal and default mode networks and the visual system (Jednoróg et al., 2012; Mackey et al., 2015). One study focused only on the PFC also found similar reductions in thickness (Lawson et al., 2013). Broadly, these regions are involved in a wide range of cognitive processing including working memory, cognitive control, autobiographical memory, theory of mind, and visual processing (Cole & Schneider, 2007; Corbetta, Kincade, & Shulman, 2002; DiCarlo, Zoccolan, & Rust, 2012; Spreng & Grady, 2010). Given that SES-related differences in many of these domains are well-established (Noble, McCandliss, & Farah, 2007), these findings could represent a neural mechanism explaining these SES-related differences in cognitive function. In contrast, two studies spanning a large age range did not find SES-related differences in thickness (Noble et al., 2015; Rosen et al., 2018). This could be because SES associations with cortical thickness vary across childhood and adolescence. Indeed, using the same sample as Noble and colleagues, Piccolo and colleagues (2016) found an SES by age interaction for average cortical thickness such that lower SES was associated with a more rapid age-related decrease in cortical thinning early in development while higher SES was associated with a less steep linear decline in thickness from childhood to adolescence. These findings are consistent with the hypothesis that low SES is associated with accelerated maturation of the cortex.

Linear decreases in cortical thickness from infancy to adulthood are well-established (LeWinn et al., 2017; Vijayakumar et al., 2016; Wierenga et al., 2014), although the mechanisms by which this pattern emerges remain in question. One interpretation of is that synaptic connections that are underutilized or inefficient are pruned, allowing the brain to adapt to the environment in which it develops (Huttenlocher, 1979; Petanjek et al., 2011; Rakic, Bourgeois, Eckenhoff, Zecevic, & Goldman-Rakic, 1986). If pruning is the primary mechanism driving cortical thinning, it is possible that ELA-related differences in cortical thickness are due to accelerated pruning. In the case of deprivation-related experiences, this may be due to a lack of experience with socially or cognitively stimulating environments (McLaughlin, Sheridan, & Nelson, 2017). Alternatively, greater pruning could reflect precocious specialization and maturation of circuits utilized more frequently by children exposed to ELA; in the absence of behavioral data associated with specific patterns of cortical thinning, caution is warranted in interpreting these patterns as either adaptive or maladaptive (ME., MD., & IH., 2016). Other work suggests that age-related decreases in cortical thinning may actually be due to increases in myelination across development (Natu et al., 2018; Sowell et al., 2004). Increased myelination, which is most pronounced in deeper cortical layers may increase the intensity of voxels at the grey-white matter border, therefore making the cortex appear thinner across age. If myelination is the primary mechanism by which cortical thinning happens, it is possible that increased cortical thickness in response to ELA may be due to faster development of structural connectivity between regions. Of course, these mechanisms are not mutually exclusive and future longitudinal work measuring multiple forms of ELA utilizing both T1-weighted imaging and diffusion tensor imaging is needed to disentangle the precise mechanisms by which ELA leads to thinner cortex in youths.

#### Amygdala-PFC Connectivity

Existing work examining ELA and task-related amygdala-PFC connectivity has produced mixed findings. Across both threat and deprivation, approximately half of the studies observed that ELA was associated with more negative functional connectivity, indicating accelerated development (Colich et al., 2017; Gee, Gabard-Durnam, et al., 2013; Keding & Herringa, 2016), while several others showed the opposite pattern of results such that youths exposed to ELA demonstrate more positive amygdala-PFC connectivity than non-exposed youths, indicating delayed development (Cisler, James, et al., 2013; Marusak et al., 2015; Silvers, Insel, et al., 2016a). Therefore, existing work has yet to provide clear evidence for an association between ELA and accelerated development of these systems. Moreover, there is no clear evidence that specific types of adversity have differential influences on the development of this circuit.

One possibility is that amygdala-PFC functional connectivity is not a reliable marker of neural development. Unlike cortical thickness which has been studied widely across large representative samples (for review see Vijayakumar et al., 2016), research documenting amygdala-PFC connectivity as a marker of maturation is more modest (Gee, Humphreys, et al., 2013; Kujawa et al., 2016; Silvers, Insel, et al., 2016b; Wu et al., 2016), and to our knowledge, all of the studies that have demonstrated a developmental shift in this circuit have been cross-sectional. As such, amygdala-mPFC connectivity may be an unreliable marker of neural maturation. Alternatively, while all these tasks focused on some sort of emotional processing, it is possible that heterogeneity across different tasks may contribute to differences in results. Future longitudinal work with a range of emotional processing tasks will be needed to establish the developmental trajectory of amygdala-mPFC connectivity to determine whether it is a robust metric of development.

### Effect of ELA Across Multiple Domains of Biological Aging

Given the range in operationalizing accelerated development and potential mechanisms linking ELA and accelerated biological development, it is surprising that few have attempted to reconcile across these different metrics of maturation. Only three studies to our knowledge have incorporated multiple metrics of accelerated development in adolescence. Belsky and Shalev (2016) put forth a “two-hit” model suggesting that ELA accelerates development first through telomere erosion and second, through earlier reproduction, which can increase oxidative stress and accelerate telomere erosion. Although this model accounts for two forms of accelerated aging, it does not directly compare and contrast the effects of ELA on both metrics of accelerated development – cellular aging and pubertal timing. Sumner and colleagues (Sumner et al., 2019) examined how exposure to threat and deprivation-related ELA influenced both DNAm age and pubertal timing. They found that exposure to threat, but not deprivation, contributes to both accelerated DNAm age and accelerated pubertal timing. Understanding the effect of ELA across domains of biological aging is important as it reflects a unique or shared mechanism linking ELA and accelerated development. There is also evidence to suggest that pubertal timing and cellular aging are highly correlated (Binder et al., 2018), suggesting a potential shared mechanism contributing to the development of both domains. However, other work demonstrates variation in the rate of change across different metrics of biological aging (D.W. Belsky et al., 2015), indicating that multiple mechanisms might underlie the ELA-accelerated development association, depending upon the metric of accelerated development. Although increased allostatic load has been proposed as a mechanism linking ELA to accelerated pubertal timing (Danese & McEwen, 2012; McEwen, 1998), empirical evidence testing this possibility is currently lacking. Moreover, allostatic load is a multi-dimensional construct involving numerous biological systems, and it is unclear if accelerated weathering occurs across all systems to a similar degree (Geronimus, 1992; Geronimus, Hicken, Keene, & Bound, 2006). If allostatic load is a mechanism contributing to accelerated pubertal development, it could explain our disparate findings regarding threat and deprivation as exposure to early-life trauma has been consistently associated with elevated allostatic load (Danese & McEwen, 2012; Scheuer et al., 2018; Widom, Horan, & Brzustowicz, 2015), whereas recent work indicates that even extreme exposure to deprivation association with institutional rearing is unrelated to allostatic load (Slopen et al., 2019). It is clear that greater work is needed to elucidate the mechanisms underlying accelerated development following exposure to ELA across domains, whether they are global or specific to particular dimensions of early experience, and how these mechanisms ultimately contribute to changes in reproductive function, cellular aging, and brain development.

### Implications of Accelerated Development

Accelerated aging across domains has been associated with a host of mental and physical health problems. For instance, accelerated pubertal timing is linked with a range of mental health problems including heightened levels of risk-taking behavior, delinquency and substance abuse problems (Copeland et al., 2013; Harden & Mendle, 2012), as well as depression and anxiety disorders (Hamilton, Hamlat, Stange, Abramson, & Alloy, 2014; Mendle, Harden, Brooks-Gunn, & Graber, 2010; Mendle et al., 2014; Negriff & Susman, 2011; Ullsperger & Nikolas, 2017). Accelerated pubertal timing is also associated with a range of physical health problems, including cardiovascular disease, polycystic ovarian syndrome in females, and testicular cancer in males (Day, Elks, Murray, Ong, & Perry, 2015; Golub et al., 2008; Lakshman et al., 2009; Velie, Nechuta, & Osuch, 2006). Accelerated cellular aging has also been associated with depression (Ridout, Ridout, Price, Sen, & Tyrka, 2016), anxiety (Malouff & Schutte, 2017), posttraumatic stress disorder (Li, Wang, Zhou, Huang, & Li, 2017), cardiovascular disease (Rehkopf et al., 2016), cancer (Zhu et al., 2016), and all-cause mortality (Needham, Rehkopf, et al., 2015). Finally, altered trajectories of cortical development have been linked to variations in general intelligence (Shaw et al., 2006) and attention-deficit hyperactivity disorder (McLaughlin, Sheridan, Winter, et al., 2014). Little research has directly examined whether accelerated development in these systems is a consequence of pre-existing mental and physical disorders, or a mechanism explaining elevated risk for mental and physical health problems in youth who have experienced ELA (see J. Belsky, Ruttle, Boyce, Armstrong, & Essex, 2015; Colich et al., 2019; Mendle et al., 2014; Negriff, Saxbe, et al., 2015 for work that has explored this idea). For instance, some evidence suggests that accelerated pubertal timing explains a significant proportion of the association between threat-related ELA risk for mental health problems in adolescence (Colich et al., 2019), and that telomere shortening occurs prior to the onset of depression in an at-risk population (Gotlib et al., 2015). However, there is also evidence to suggest that early psychosocial difficulties precede early pubertal onset (Mensah et al., 2013) and could potentially accelerate cellular aging as well (Lindqvist et al., 2015). A key issue for future research will be to determine whether early interventions targeting psychosocial mechanisms linking ELA with mental and physical health problems are capable of altering observed patterns of accelerated biological aging.

It is also important to acknowledge that although there are strong associations among accelerated development and negative mental and physical health outcomes, accelerated development is most likely an adaptation to current and presumably future environmental conditions (J. Belsky, 2019). In a highly dangerous or unpredictable environment, it may be adaptive in the short-term to reach adult-like capabilities at an earlier age, in order to either reach reproductive status earlier, or reach independence from the caregiving situation at an earlier age. This immediate goal may outweigh the longer-term consequences of mental and physical health problems. If the environment is signaling imminent mortality, then this trade-off is one that is evolutionarily adaptive. It will be important that future work consider the adaptive significance of accelerated development in response to ELA.

### Limitations and Future Directions

Several limitations of this work highlight key directions for future research. First, we examined a relatively small number of studies for some domains and within each dimension of adversity, particularly with regard to cellular aging and brain development. More research is needed to evaluate whether all forms of adversity influence cellular aging or whether these associations are stronger for experiences of threat. Similarly, due to the small number of studies published on these variables, we collapsed across measures of cellular aging, including telomere length and DNAm age. These markers reflect distinct biological processes with differing molecular signatures, and we recognize that combining across these two metrics of cellular aging is most likely an over-simplification of the effects of ELA on cellular aging. These findings should be replicated when more studies have been published on the associations of ELA with both telomere length and DNAm age. Similarly, it is important to note that we did not include studies examining the effects of ELA on methylation patterns of single genes due difficulties in understanding what typical developmental patterns of specific gene methylation would be. For a systematic review of the effects of ELA on gene-specific methylation patterns see Lang et al. (2019). Second, although we show associations among ELA, pubertal timing, cellular aging, and cortical thinning, we do not have the data to speak to the underlying mechanisms driving these associations. Although we speculate that accelerated biological development following ELA is most likely due to the effects of ELA on allostatic load, future experimental work should investigate the underlying biological mechanisms supporting these associations. Finally, in only examining samples in childhood and adolescence, we hoped to examine the associations between ELA and accelerated development, independent of psychopathology. However, given strong links among ELA, accelerated development and psychopathology, it is impossible to confidently conclude that psychopathology is not driving the effects of ELA on accelerated biological development. Future longitudinal work using at-risk samples should address the directionality of these associations to determine with confidence the direct effect of ELA on accelerated biological aging.

### CONCLUSIONS

Through meta-analysis and systematic review, we support the idea that ELA accelerates biological aging, as measured by pubertal timing, cellular aging, and cortical thinning in childhood and adolescence. However, these associations varied systematically as a function of adversity type. Specifically ELA characterized by threat, but not deprivation or SES, was associated with accelerated pubertal development, suggesting specificity in the link between ELA and pubertal timing. ELA was also associated with accelerated cellular aging as measured by both leukocyte telomere length and DNA methylation age, with the strongest evidence for threat-related ELA being associated with cellular aging. ELA was consistently associated with accelerated cortical thinning, with threat-related ELA associated with ventromedial PFC thinning and deprivation and SES more consistently associated with thinning in the frontoparietal, default mode, and visual networks. We found no consistent association of ELA with amygdala-mPFC functional connectivity. These findings suggest both common and specific associations of dimensions of ELA with multiple domains of biological aging and highlight the importance of delineating the mechanisms through which specific types of early environmental experiences influence different aspects of biological aging in childhood and adolescence and determining how these pathways ultimately contribute to health disparities.

