## Supplement for "Biological Aging in Childhood and Adolescence Following Experiences of Threat and Deprivation: A Systematic Review and Meta-Analysis"

Supplementary Materials

**PubMed Search Terms**

(“Violence"[Mesh] OR "Psychological Trauma"[Mesh] OR "Adoption"[Mesh] OR "Socioeconomic Factors"[Mesh] OR "Health Status Disparities"[Mesh] OR violence[tw] OR trauma[tw] OR neglect[tw] OR negligence[tw] OR maltreatment[tw] OR institutional rearing[tw] OR deprivation[tw] OR socioeconomic status[tw] OR poverty[tw] OR "early adversity"[tw] OR "early life stress"[tw]) AND ("Puberty"[Mesh] OR "Cell Aging"[Mesh] OR "Methylation"[Mesh] OR "epigenesis, genetic"[mesh] OR pubertal timing[tw] OR pubertal onset[tw] OR "onset of puberty"[tw] OR menarche[tw] OR pubertal timing[tw] OR telomere length[tw] OR methylation[tw] OR neural[tw]) AND ("infant"[mesh] OR "child"[mesh] OR "adolescent"[mesh] OR infant*[tw] OR infancy[tw] OR child*[tw] OR adolesc*[tw] OR teen*[tw] OR schoolchild*[tw] OR “pediatrics”[mesh] OR pediatric*[tw])

**Scopus Search Terms**

( TITLE-ABS-KEY ( violence  OR  trauma  OR  adoption  OR  socioeconomic  OR  deprivation  OR  neglect  OR  negligence  OR  "institutional rearing"  OR  poverty  OR  "early life adversity"  OR  "early life stress" )  AND  TITLE-ABS-KEY ( puberty  OR  menarche )  AND  TITLE-ABS-KEY ( child*  OR  adolesc*  OR  infan*  OR  pediatric*  OR  teen* ) )

( TITLE-ABS-KEY ( violence  OR  trauma  OR  adoption  OR  socioeconomic  OR  deprivation  OR  neglect  OR  negligence  OR  "institutional rearing"  OR  poverty  OR  "early life adversity"  OR  "early life stress" )  AND  TITLE-ABS-KEY ( methylation  OR  "cell aging"  OR  "telomere length"  OR  epigenetic* )  AND  TITLE-ABS-KEY ( child*  OR  adolesc*  OR  infan*  OR  pediatric*  OR  teen* ) )

( TITLE-ABS-KEY ( violence  OR  trauma  OR  adoption  OR  socioeconomic  OR  deprivation  OR  neglect  OR  negligence  OR  "institutional rearing"  OR  poverty  OR  "early life adversity"  OR  "early life stress" )  AND  TITLE-ABS-KEY ( neural )  AND  TITLE-ABS-KEY ( child*  OR  adolesc*  OR  infan*  OR  pediatric*  OR  teen* ) )

**PsycINFO Search Terms**

ti(violences OR trauma OR adoption OR socioeconomic OR deprivation OR neglect OR negligence OR institutional rearing OR poverty OR early life adversity OR early life stress) AND (puberty OR methylation OR cell aging OR telomere length OR menarche OR epigenetic* OR neural) AND (child* OR adolsc* OR infan* OR pediatric* OR teen*)

**Web of Science Search Terms**

TS=(violence OR trauma OR adoption OR socioeconomic OR deprivation OR neglect OR negligence OR "institutional rearing" OR poverty OR "early life adversity" OR "early life stress") AND TS=(puberty OR methylation OR "cell aging" OR "telomere length" OR menarche OR epigenetic* OR neural) AND TS=(child* OR infant* OR adolesc* OR teen* OR pediatric*)

**Google Scholar Search Terms**

(violence|trauma|adoption|socioeconomic|neglect|negligence|"institutional rearing"|poverty|"early life adversity"|"early life stress") AND (puberty|methylation|"cell aging"|"telomere length"|menarche|epigenetic|neural) AND (child|infant|adolescent)
